# Environmental DNA reveals invasive crayfish microbial associates and ecosystem-wide biodiversity before and after eradication

**DOI:** 10.1101/2022.05.20.492886

**Authors:** Kimberly M. Ballare, Anna Worth, Tyler Goodearly, Dannise V. Ruiz-Ramos, Eric Beraut, Hailey Nava, Colin Fairbairn, Robert K. Wayne, Beth Shapiro, Ginny Short, Rachel S. Meyer

**Author notes:** **Author Contributions:** K.M.B., G.S., T.G., and R.S.M. conceived and designed experiments and sampling scheme. G.S. coordinated site access and supervised field work. T.G. maintained and sampled aquaria. K.M.B., G.S., H.N., D.V.R-R., T.G., and R.S.M collected field samples and coordinated volunteer field sampling. K.M.B., A.W., D.V.R-R, E.B., H.N., and C.F. conducted molecular laboratory work. K.M.B., A.W., H.N., C.F, and R.S.M. conducted literature review. K.M.B. and R.S.M. curated, analyzed, and interpreted data. K.M.B. and T.G. generated the map figure and K.M.B. generated the data figures. R.K.W., B.S., G.S., and R.S.M. provided supervision, funding, materials, and analysis tools. K.M.B. and R.S.M. wrote the manuscript with input from all co-authors. **Corresponding Author:** Rachel Meyer, 1-206-351-7997.

## Abstract

Biodiversity monitoring in conservation projects is essential to understand environmental status and recovery. However, traditional field surveys can be expensive, time-consuming, biased towards visual detection, and focused on measuring a limited set of taxa. Environmental DNA (eDNA) methods provide a new approach to biodiversity monitoring that has the potential to sample a taxonomically broader set of organisms with similar effort, but many of these approaches are still in the early stages of development and testing. Here, we use multilocus eDNA metabarcoding to understand how the removal of invasive red swamp crayfish impacts local biodiversity of a desert oasis ecosystem, as well as to detect crayfish both directly and indirectly. We tracked crayfish DNA signatures, microbial DNA associated with crayfish, and biodiversity of plant, fungal, animal, and bacterial communities through time. We were unsuccessful in detecting crayfish directly in either control tanks or oases using targeted metabarcoding primers for invertebrates and eukaryotes, similar to previous studies which have shown variable levels of success in detecting crayfish from environmental samples. However, we were successful in discerning a suite of 90 crayfish-associated taxa to serve as candidate bioindicators of invasive presence using 16S and Fungal ITS2 metabarcoding. Ranking these 90 taxa by their geographic distribution in eDNA surveys and by evidence of crayfish-associations in the literature, we support 9 taxa to be high-ranking, and suggest they be prioritized in future biomonitoring. Biodiversity analyses from five metabarcode loci including plants, animals, and both prokaryotic and eukaryotic microbes showed that communities differed but that species richness remained relatively similar between oases through time. Our results reveal that, while there are limitations of eDNA approaches to detect crayfish and other invasive species, microbial bioindicators offer a largely untapped biomonitoring opportunity for invasive species management, adding a valuable resource to a conservation manager’s toolkit.

## 1. Introduction

Environmental DNA (eDNA) is an increasingly popular tool for biodiversity monitoring across landscapes and taxa. Using eDNA, reliable biodiversity surveys can be quickly and easily obtained by land managers and community scientists without need for expertise in field methods or taxonomy (Meyer et al., 2021; Suarez-Menendez et al., 2020). These approaches are particularly promising for invasive species monitoring and conservation projects (Ficetola et al., 2008; Sepulveda et al., 2020) because they make it possible for managers to simultaneously monitor biodiversity and detect invasive species using a single technique. Biomonitoring during restoration and conservation management projects is also an essential part of understanding whether restoration interventions have been successful, e.g. whether taxa of interest are present in an ecosystem. Both traditional and eDNA surveys can be useful for direct invasive species detection and monitoring, as long as the taxon in question can be reliably detected (e.g. Crane et al., 2021; Halstead et al., 2017; Ratsch et al., 2020) However, some taxa are challenging to detect and quantify using either method even when they are known to be present: including cryptic or elusive species, species persisting at low densities, or with variable detection during their life cycles.

One set of taxa that can be challenging to detect with either traditional or molecular approaches are crayfish (Astocoidea L.). When abundant, crayfish are easily trapped or detected with visual surveys (e.g. Short, 2021), but crayfish traps also risk mortality of non-target organisms (Tréguier et al., 2014). When rare, crayfish can escape detection through visual surveys or trapping because they can burrow in sediment and find refugia for up to several months (Pritchard et al., 2021), potentially leading to false negative detection in early stages of invasion, or false successes of eradication programs for invasive species (e.g. Nunes et al., 2017). Molecular detection approaches also have variable levels of success (Chucholl et al., 2021; Dougherty et al., 2016; Harper et al., 2018; Ikeda et al., 2016; Troth et al., 2020). For the globally invasive red swamp crayfish (*Procambarus clarkii* Giard, 1852) in particular, targeted detection assays from environmental samples have not proven to be a stable monitoring approach (Cai et al., 2017; Mauvisseau et al., 2018; Thalinger et al., 2021). For example, in a 2014 study, *P. clarkii* was not reliably detected from environmental samples even when using species-specific primers tested and designed for the study (Tréguier et al., 2014). Additionally, some assays fail when researchers modify protocols such as changing filters, extraction volumes, and substrates (Geerts et al., 2018; Tréguier et al., 2014), or when attempting to detect crayfish at different points in their life cycle (Dunn et al., 2017). One potential explanation is that crayfish do not shed DNA reliably into the environment (Curtis and Larson, 2020), which is also a known issue with detecting other crustacean and invertebrate species in marine or aquatic ecosystems (e.g. green crab, Crane et al., 2021).

To mitigate problems with direct detection of a particular species, an emerging eDNA method is to use microbial proxies as bioindicators of sentinel species for tracking invasion and other disturbances (Astudillo-García et al., 2019). The characterization of microbial communities can also provide vital information about the biology of the target species (Ficetola et al., 2019), as well as characterize the community ecology of the site. Microbial proxies used for indirect detection can include both infectious agents (Wittwer et al., 2018), as well as more benign associates such as gut microbes, diet species, and symbionts (Ficetola et al., 2019; Skelton et al., 2017). While the use of eDNA metabarcoding is appealing because it can detect many species simultaneously and be used on a variety of substrates (Jarman et al., 2018), few studies use metabarcoding profiles to generate bioindicator proxies for target species detection. It may be difficult to detect “true” microbial bioindicators, that is, a taxon may appear driven by a target species, but in fact that microbe is common in many environments and has broad ecological associations and roles. However, because *P. clarkii* and other crayfish have multiple known symbionts, pathogens, and other microbial associates verified across many studies (Chen et al., 2021; Dragičević et al., 2021), this makes it an ideal taxon to test detection success through microbial proxies. Because of the commonness of invasive red swamp crayfish across the globe (Geiger et al., 2005), the outsized negative impacts that they have on local biodiversity and ecosystem function (Oficialdegui et al., 2020; Souty-Grosset et al., 2016), and the known detection challenges, multiple methods of detection are clearly needed.

Here, we explore how environmental DNA can be used to track the invasive red swamp crayfish, as well as reveal novel components of the biology of the invaded ecosystem. We tested multiple existing published assays for direct detection of *P. clarkii*, and used universal metabarcoding to 1) detect signals of crayfish presence, 2) generate a reliable list of microbial crayfish bioindicators, and 3) track the biotic effects of crayfish presence and pesticide eradication methods on the overall ecological community. Because the red swamp crayfish has such a large known effect on aquatic biodiversity, consuming living plants, detritus, and animals as an opportunistic omnivore (Alcorlo et al., 2004; Bucciarelli et al., 2019), we expect to see discernable correlations with the surrounding biotic community including changes in community composition and species richness over time.

## 2. Methods

### 2.1 Study System

We conducted our study in two palm oases in the Thousand Palms Oasis Preserve (Thousand Palms Oasis and Simone Pond, Figure 1) managed by the Center for Natural Lands Management (CNLM), as well as in an experimental tank system (see section 2.3 below). Located in the Colorado Desert, California, USA, the Thousand Palms Oasis Preserve contains numerous distinct oases fed by a common underground aquifer, with the vegetative community dominated by the native California Fan Palm (*Washingtonia filifera)*. Simone Pond is distinct from other oases in the preserve as it is characterized by an open habitat and more consistent water depth throughout the year. Thousand Palms Oasis is located approximately one km south of Simone Pond. Surrounded by the same dominant vegetation community, it differs from Simone Pond in that it has less light that permeates through the fan palm canopy, and a shallow water depth. Other than Western mosquitofish *Gambusia affinis* which is an invasive species that persists in both oases, Thousand Palms Oasis has no fish species. There is no record of crayfish existing in Thousand Palms Oasis.

**Figure 1.**
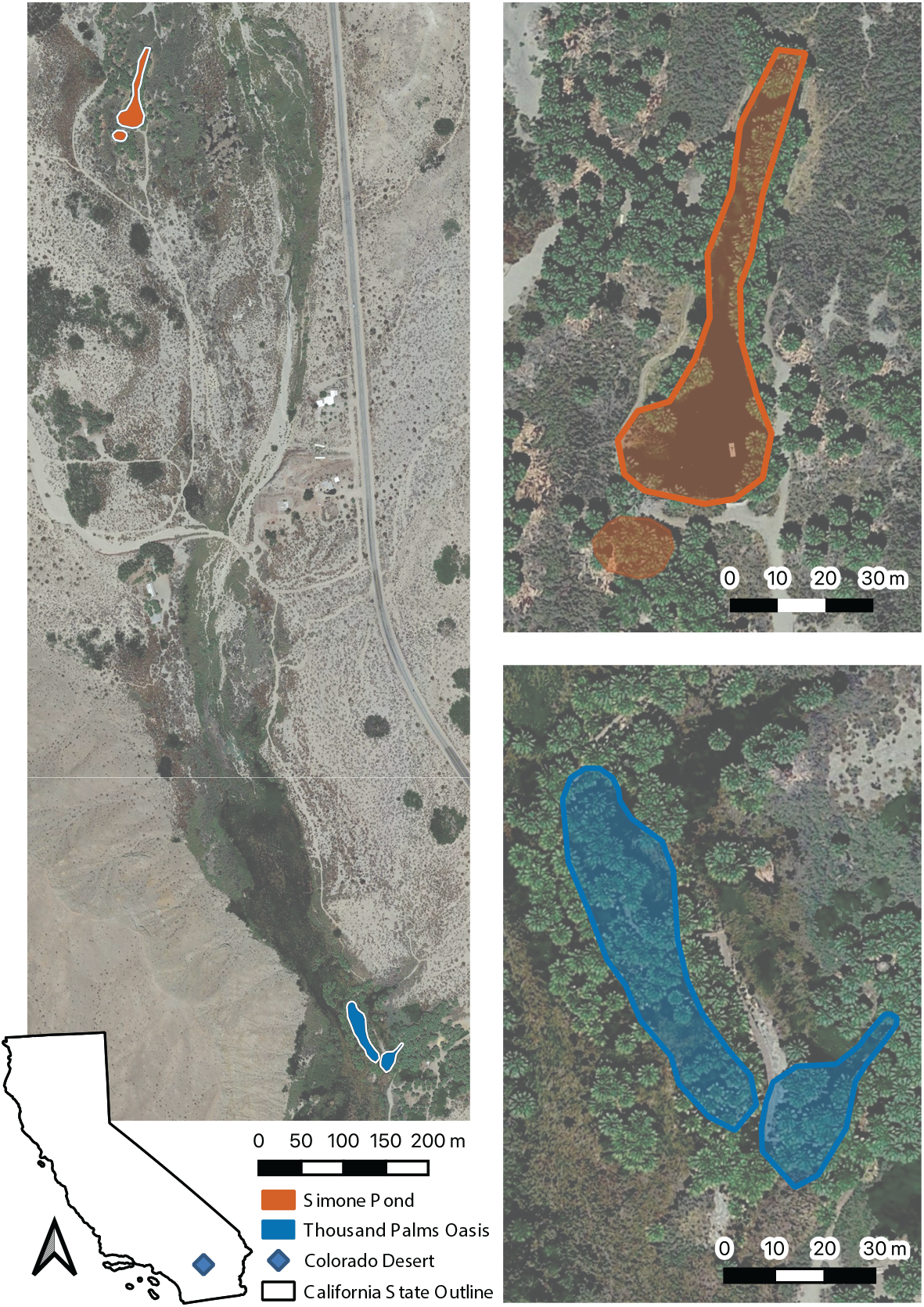
Map of study area, located in Colorado desert, Southern California, USA. Left, Aerial view of Thousand Palms Oasis Preserve. Top right, Simone Pond, oasis invaded with *Procambarus clarkii*. Bottom right, Thousand Palms Oasis, not invaded. Shaded portions indicate inundated areas.

Simone Pond has had multiple invasive species introductions, including red swamp crayfish (*Procambarus clarkii)* in the 1950’s, and tilapia (*Oreochromis* sp.) in 2015, both of which caused significant biodiversity loss in Simone Pond (Short, 2021). Importantly, Simone Pond has unique geophysical characteristics including an open habitat that allows for light permeation, high water quality, and submergent and emergent vegetation, which make it a candidate location for the introduction of the rare and endangered desert pupfish (*Cyprinodon macularius* Baird & Girard 1853, Martin and Saiki, 2005; Varela-Romero et al., 2002). Unfortunately, previous attempts to introduce desert pupfish to Simone Pond were unsuccessful due to competition with and predation by red swamp crayfish. CNLM Staff used several methods to try to remove the invasive species and found cypermethrin (Cyper TC®, Control Solutions Inc., EPA Reg. No. 53883–92 [(±)α-cyano-(3-phenoxyphenyl)methyl(±) - cis/trans-3-(2,2- dichloroethenyl)- 2,2-dimethylcyclopropanecarboxylate]) to be the most effective at controlling crayfish (Short, 2021).

Cypermethrin is an insecticide that kills target organisms by blocking voltage-gated sodium channels to affect the peripheral nervous system (Shafer et al., 2005). Derived from the insecticidal compounds found in chrysanthemum, cypermethrin is known be more toxic to crustaceans than vertebrate fish species (Morolli et al., 2006; Sandodden and Johnsen, 2010; Soares et al., 2019), with mild toxicity present in amphibians and mammals (Edwards et al., 1986; Khan et al., 2009). Cypermethrin and related compounds have been shown to be effective at eradicating invasive crayfish populations at low doses (e.g. Sandodden and Johnsen, 2010), although it is also toxic to other aquatic organisms including benthic invertebrates (Zhang et al., 2018). Prior to applying the pesticide, preserve staff drained water to a temporary reservoir, and manually filtered all visible tilapia and crayfish. After the initial cypermethrin application to the drained Simone Pond, they allowed Simone Pond to refill naturally. Pesticide application in Simone Pond began in July 2019 and continued monthly until July 2020 (Pesticide research authorization permit #1905023, California Department of Pesticide Regulation). During and after the application of the pesticide CNLM staff conducted regular monitoring including trapping and visual surveys, which revealed a substantial population of crayfish in Simone Pond. Prior to pesticide application, over 6,300 crayfish were removed via baited minnow traps between January and June 2019, with an estimated 26,000 individuals present in January 2019 (Leslie Depletion Model; Leslie and Davis, 1939). No crayfish or tilapia were observed for 100 days after initial application of the pesticide, but one dead crayfish was found 101 days after the initial application. As of May 2022, no others have been observed since October 2019 (Short, 2021).

### 2.2 Environmental Sample Collection

We used environmental DNA sampling to detect crayfish and monitor changes in biodiversity in Simone Pond and Thousand Palms Oasis before, during, and after the cypermethrin application program. Specifically, we collected environmental samples at four time points (Table 1, Supplemental Table S1): 1) June 2019, immediately before cypermethrin application began; 2) December 2019, after 5 monthly applications of cypermethrin; 3) June 2020, after the last monthly cypermethrin application; and 4) Dec 2020, 5 months after the last application of cypermethrin. We collected sediment (fully water submerged to rarely water submerged surfaces of the oasis, containing soil usually mixed with algal and siliceous matter) and soil (terrestrial surface, never water submerged) samples during each sampling event, and water samples during the June 2019 sampling event. Water samples were collected using a randomized transect approach in each oasis. One liter of water was collected in an IV fluid bag and passively drip filtered through a 0.22 um Sterivex filter (Millipore, Burlington, Massachusetts, USA). Filters were immediately refrigerated, and DNA was extracted the following day. Sediment or soil samples were collected at regular spatial intervals around the water edge of the oasis, with sampling locations selected so as to maximize environmental heterogeneity by taking samples from sites that were rarely, frequently, or fully submerged, from under palm trees, in aquatic grass beds, in dense algal masses, and along oasis perimeter fencing. At each site, we collected a sample of three 2ml cryotubes filled with soil or sediment. We obtained these substrates by scraping off the surface with the tube itself (soil) or by obtaining an underwater surface sediment sample using a 2-inch diameter PVC pipe and scraping sediment off of it with the tube itself. Water and substrate samples were immediately frozen at −20°C and transferred to −80°C for archiving. Immediately before DNA extraction, we thawed soils and sediments on ice, homogenized them by vortexing, and pooled an aliquot from each tube together to represent the single site sample. This mixture was again homogenized and 0.25g was taken for DNA extraction.

**Table 1.** Sampling Summary

| Sample Date | Simone Pond Condition | N Samples Simone Pond |  |  |  | N Samples Thousand Palms Oasis |  |  |  | N Samples Tanks |
| --- | --- | --- | --- | --- | --- | --- | --- | --- | --- | --- |
|  |  | Water | Sediment | Soil | Total | Water | Sediment | Soil | Total | Water Only |
| Jun 2019 | Invasives Present | 4 | 21 | 0 | 25 | 10 | 27 | 6 | 43 | 13 |
| Dec 2019 | Cypermethrin applied monthly since July 2019 | 0 | 31 | 3 | 34 | 0 | 14 | 2 | 16 | 0 |
| Jun 2020 | Cypermethrin applied monthly since July 2019 | 0 | 31 | 6 | 37 | 0 | 15 | 5 | 20 | 0 |
| Dec 2020 | No mitigation for 5 months | 0 | 28 | 3 | 31 | 0 | 8 | 8 | 16 | 0 |

### 2.3. Aquarium eDNA Sampling

In June 2019, we established thirteen four-gallon aquarium tanks using unfiltered well water from the preserve source aquifer. The aquaria contained four treatments: four crayfish-only tanks (three containing live crayfish, and one with a crayfish carcass), three tilapia-only tanks, three crayfish and tilapia tanks, and three no-animal (blank) tanks. Crayfish were not fed for the duration of their time in the tanks to avoid introducing a foreign food microbiome. After two weeks, we collected a single one-liter water sample from each tank and processed samples for DNA extraction as described in section 2.2 above. Non-invasive sampling of these tanks was compliant with NRC guidelines of animal treatment, and was an approved component of CNLM Project G2023 (Short, 2021).

### 2.4. DNA Extraction, Library Preparation, and Sequencing

#### 2.4.1 Metabarcoding

We extracted soil and sediment samples using a standard 0.25g input into the Qiagen DNeasy PowerSoil Kit (Qiagen Inc., Germantown, MD) and followed manufacturer’s instructions. We extracted water DNA from the Sterivex filter cartridge using the Qiagen DNeasy Blood and Tissue Kit following the protocol reported in Spens et al. (2017). From soil, sediment, and water DNA extracts, we amplified 5 metabarcode markers to detect different taxonomic groups: 16S targeting Bacteria and Archaea (515F 806R; Caporaso et al., 2012), 18S targeting Eukaryota (Euk_1391f and EukBr; Amaral-Zettler et al., 2009), fungal ITS1 targeting Fungi (abbreviated hereafter as “FITS”, ITS5 and 5.8S; White et al., 1990) and plant ITS2, targeting plants and green algae (abbreviated hereafter as ‘PITS’, ITS-S2F and ITS-S3R; Gu et al., 2013), and CO1, initially targeting the full region (LCO1490 and HCO219; Hebert et al., 2003) to search for crayfish DNA in the June 2019 sample. After preliminary sequence data indicated high proteobacteria swamping in the initial CO1 samples, we switched to the partial CO1 primers for subsequent samples targeting both invertebrates and protists (mlCOIintF and Fol-degen-rev; Leray et al., 2013).

For each marker, we performed three replicate PCRs to account for stochastic amplification. We then pooled PCR replicates by marker, quantified the amount of DNA in each pool using the Qubit ™ dsDNA Broad Range assay (Invitrogen™), and finally pooled all reactions per sample of five markers together using equimolar concentrations of each reaction. We indexed the amplicons in each sample pool using dual unique Nextera indices (UMI), then combined all metabarcoding libraries for sequencing on either an Illumina MiSeq with the v3 kit for 2×300bp reads (18S, CO1, PITS (all sampling points), and 16S and FITS for the June 2019 samples) or a NextSeq 550 for 2×150bp reads (for 16S and FITS in Dec 2019, June 2020, and Dec 2020). Sequencing was conducted at UCLA or UCSC sequencing facilities to produce fastq files of DNA libraries.

To provide Preserve staff with preliminary results after each sampling event we processed and sequenced each of the four sample batches separately. To test for batch effects, we randomly selected 10 samples from each sample date (five samples per oasis), re-extracted DNA, amplified two markers (16S and 18S), and sequenced them together on the NextSeq as described above. We co-analyzed these with the original samples to detect whether community composition of replicates fell within their respective batch groups in ordinations and permutational ANOVA.

We analyzed metabarcoding sequence data using the *Anacapa Toolkit* (Curd et al., 2019), which uses CutAdapt (Martin, 2011) and FastX-Toolkit (Gordon and Hannon, 2010) for Fastq quality control and filtering, and DADA2 (Callahan et al., 2016) for amplicon sequence variant (ASV) assignment and chimera detection and removal. Taxonomic assignment for each ASV was made using Bowtie2 (Langmead and Salzberg, 2012) and the Bayesian Lowest Common Ancestor (BLCA; Gao et al., 2017) algorithm with our metabarcode-specific CRUX reference databases (version 2; made from October 2019 NCBI data; doi:10.5068/D1QX0W), of which the 18S and CO1 databases included sequences from *P. clarkii*. We assumed the default parameters except the minimum quality score was set to 32. Taxonomic assignments at each classification level were retained when the bootstrap confidence cutoff score was over 70 for 16S and FITS or over 80 for CO1, 18S, and PITS. The ASVs with the same resulting taxonomic entry from domain to species were summed to be a single entry as the species-equivalent. We converted metadata and ASV tables (Supplemental Table S1) to *Phyloseq* (McMurdie and Holmes, 2012) objects for analysis using *Ranacapa* (Kandlikar et al., 2018). We filtered data for analysis first by decontaminating tables with the decontam package (Davis et al., 2018) and prevalence set to 0.1, and then removing any taxa represented by fewer than 10 total reads across all samples.

#### 2.4.2 Metagenome

After cypermethrin crayfish control began in Simone Pond, hundreds to thousands of killed crayfish carcasses degraded in Simone Pond, and pieces of the exoskeletons were occasionally found intact. In December 2019, we found a piece of crayfish carapace exoskeleton in the Simone Pond surface sediment. To examine the metagenome signatures on this exoskeleton and test for any retained crayfish DNA in decomposing carcasses, we washed the sample with pure water and ground 0.25g with a Qiagen PowerLyzer Homogenizer and extracted DNA. A Tn5 library was prepared according to Russell et al. (2020) and sequenced using an Illumina NextSeq 550 (Illumina, San Diego, California). We used bbduk from BBTools (Bushnell, 2014) to trim low quality (q<25) reads and remove adapters. Then, we merged reads with bbmerge and mapped them to the *P. clarkii* genome (ASM2042438v2) using BWA (v 0.7.17-r1188; Li and Durbin, 2009). Low entropy mapped reads were removed using bbduk with entropy=0.9 entropywindow=25 entropyk=10 entropymask=f. Merged reads were matched to taxa through MG-RAST (Meyer et al., 2008).

#### 2.4.3 Quantitative PCR

In an attempt to have more sensitive direct detection of crayfish than metabarcoding, we also used published quantitative PCR (qPCR) assays to amplify *P. clarkii* from samples where crayfish were present. We selected two primer sets that reported reliable detection of red swamp crayfish from freshwater DNA extracts: Mauvisseau et al. (2018) [CO1-Pc-03-F 5’-GGAGTTGGAACAGGATGGACT-3’, CO1-Pc-03-R 5’-AATCTACAGATGCTCCCGCA-3’]; and Knudsen et al. (2019) [Procla_co1_F04 5’-GCGGGAGCATCTGTAGATTT-3’, Procla_co1_R04 5’-ATAGCTCCTGCCAACACAGG-3’]. The templates we screened included known crayfish positive and known crayfish negative samples from the June 2019 DNA extracts: three sediment samples and two water samples from Simone Pond (which contained an estimated 26,000 crayfish), one water sample and two sediment samples from Thousand Palms Oasis (which had no crayfish), and eight water samples from the tank experiments (5 samples from crayfish positive and 3 samples from crayfish negative tanks). Reactions were performed in triplicate and included an 18S reaction (same 18S primer as used for metabarcoding) as a positive control. Melt curve temperatures were also recorded. The published assays used different annealing temperatures: 60 C° (Knudsen et al. 2019) and 59 C° (Mauvisseau et al. 2018). Using iTaq Universal SYBR Green Supermix (BioRad, Hercules, California, USA), we ran these assays twice, once at each annealing temperature, and found results were similar, so we report the assay at 59 C°. We note we did not use FAM probes as in original publications for these assays, as the probes were solely designed to screen out other related crayfish in multi-species systems; we had no crayfish present in our system other than *P. clarkii*.

### 2.5 Statistical Analysis

#### 2.5.1 Bioindicator Analyses

We calculated microbial bioindicators of crayfish presence using community similarity percentage analysis (SIMPER; Clarke, 1993) in the R package *vegan* (Oksanen et al., 2015). SIMPER performs pairwise comparisons of species within each sample category and finds the average contributions of each species to overall Bray-Curtis dissimilarity, as well as the likelihood of each species occurring in one site type over another. From the June 2019 (pre-eradication) sample batch, we grouped together all tank and oasis samples (sediment, soil, and water) containing crayfish, and grouped together all tank and oasis samples without crayfish. We calculated the contribution of each taxon to the Bray-Curtis dissimilarity between groups containing crayfish and those without. We performed this twice: both the log (x+1) sequence read abundance and presence-absence data of sample sets rarefied to 3000 reads with 1000 permutations. To mitigate the tendency of SIMPER to preferentially detect more variable taxa as having between-group effects (Warton et al., 2012), we only considered taxa as significant drivers of community difference if they were significant in both the presence-absence and log (x+1) read abundance transformations (as per Ballare et al., 2019). We repeated the SIMPER analysis for each time point post-eradication.

We considered the taxa that were found to be significant drivers of community differences in the June 2019 SIMPER analyses to be ‘candidate bioindicators’ of crayfish presence. We vetted these candidate bioindicator taxa in multiple ways. 1) We scored geographic distribution by cross referencing each taxon within 1000 eDNA samples collected across California that used the same assay (CALeDNA publicly available data, ucedna.com; Meyer et al., 2021). Bioindicators that were present in most of these samples were given a low score because they are clearly not likely to be crayfish-specific based on distribution. Bioindicators with regional or restricted distribution, such as only in watersheds, or restricted to the oases or Coachella Valley were given high scores. 2) We scored bioindicators by information on their ecological roles and any reported associations with crayfish species. To review these bioindicator roles, we conducted a literature review using GLOBI API Plus (Poelen et al., 2014), and also reviewed literature in Google Scholar searches for the bioindicator taxon at the species, genus, or family level and ‘crayfish’. From these, we developed a composite score to and assigned each taxon a bioindicator strength weight from 0-6 (0 having no evidence based on geography or role that there is a relationship with crayfish, and 6 being tightly related to both crayfish presence based on role and specificity in our ecosystem), using the following scoring logic: 1 point for being found in under 200 samples out of the 1000 CALeDNA samples, 1 point for geographic range more specific than all of California (e.g. southern California or Coachella Valley), 2 points if most occurrences are in Simone Pond, and 2 points for having a published direct interaction with crayfish (any genus). If there was no direct crayfish interaction found in the manual literature review or GLOBI search, 1 point was added for co-occurring with crayfish or any interaction with Arthropoda. All points were taken away if found mostly in Thousand Palms Oasis.

#### 2.5.2 Ecosystem Biodiversity Analyses

We used R packages *Phyloseq* (McMurdie and Holmes, 2012) and *vegan* (Oksanen et al. 2015), for community biodiversity analyses visualized plots using *ggplot2* (Wickham et al., 2016). For these analyses we removed repeat samples and conducted tests only on sediment samples to avoid differences in community due to sample type (soil, sediment, or water), as variable compositions of clay, sand, and silt between soil and sediment samples may cause differing rates of DNA absorption and retention (Barnes et al., 2014; Cai et al., 2006). Sediment also tends to be a more reliable substrate to detect biodiversity over time as it retains a much longer record of DNA than water samples (days to weeks or even months, Turner et al., 2015; Wei et al., 2018). We examined alpha and beta diversity patterns for each metabarcode dataset separately (16S, 18S, FITS, PITS, and CO1), comparing biodiversity between oases, both within and between timepoints. To evaluate the changes in the Arthropoda community that we suspected was sensitive to cypermethrin treatment, we merged the CO1 and 18S data and subsetted it to Arthropoda only, and then conducted the same biodiversity analyses to test for differences in alpha and beta diversity of arthropods in both Simone Pond and Thousand Palms Oasis over the four time points. We conducted Analysis of Variance (ANOVA) and post-hoc Tukey tests to test for differences in observed alpha diversity over time, and permutational ANOVA (PERMANOVA) to test for significant differences in community composition at different time points in the different oases, using the Jaccard distance metric. Although PERMANOVA is more robust than other permutation methods (e.g. ANOSIM, Mantel test) to detect differences in community composition in the presence of uneven dispersion (Anderson and Walsh, 2013), we also conducted post-hoc beta dispersion tests (PERMDISP2) to test whether significant community differences detected by PERMANOVA were due to differences in dispersion rather than true biological difference.

Our analysis to look for batch effects indicated that the repeated samples did not separate from the other batches in NMDS ordination space, indicating little effect of sequencing batch on community composition (Supplemental Figure S1). However, PERMANOVA analyses of 18S and 16S data sets with the repeat samples included indicated some small but significant community differences between sample processing dates (PERMANOVA: 18S F=2.90, p<0.001; 16S F=10.864, p<0.001). To remain conservative, subsequent PERMANOVA for all markers included the processing batch as a stratum to correct for possible differences in community due to batch effects only. We visualized overall community similarity between time points and oases using NMDS ordination.

## 3. Results

### 3.1. Direct Detection of Crayfish DNA Signatures

#### 3.1.1. Metabarcoding

We expected to find crayfish DNA signatures in metabarcoding results from COI and 18S loci, but from the 84 total samples analyzed from July 2019 that included 21 sediment samples from Simone Pond (Simone Pond), 4 water samples from Simone Pond, and the 13 experiment tanks of which 7 contained crayfish, we only detected a signal of Decapoda (the order for crayfish) in 18S from 3 samples. While in all cases *Procambarus clarkii* was the most congruent sequence, it was not assigned to high confidence (ASVs required a bootstrap confidence score above 80 to be included in our final dataset). These positive samples were all from water samples: from two of the experimental tanks with crayfish and tilapia, and one from Simone Pond. Given that water retains a shorter temporal record of DNA compared to sediment, and only one natural oasis sample yielded a positive result even when an estimated ~26,000 crayfish were in the oasis, we determined metabarcoding for the purpose of direct detection would not be useful for crayfish monitoring.

#### 3.1.2. Crayfish qPCR

We found that qPCR using both sets of published *CO1* primers for *P. clarkii* were also unstable at detecting crayfish, producing false positive amplification signals from two negative controls. While some known crayfish positive samples did show qPCR amplification with both published primer sets, most failed to amplify crayfish, such as water samples from Simone Pond and the tank with the crayfish carcass (Supplemental Table S2).

#### 3.1.3. Crayfish DNA in exoskeleton material

Because our qPCR results did not detect crayfish DNA from water containing a rotting carcass, this led us to ask how much crayfish DNA persists in carcass material. Only 121 out of 602,844 merged reads mapped the carapace reads to the *Procambarus clarkii* genome (0.02%), indicating severe DNA degradation. MG-RAST metagenome analysis of the carapace reads revealed over 99% of the reads were microbial and dominated by Proteobacteria (Supplemental Table S3).

### 3.2. Bioindicator Analysis

#### 3.2.1. Candidate Crayfish Bioindicator taxa from SIMPER analysis

From the samples collected in July 2019, using 16S and FITS datasets, we detected 90 microbial taxa that were significantly more likely to occur in samples containing crayfish (crayfish tanks and Simone Pond) than samples where crayfish were known to be absent (blank tank, tilapia-only tanks, and Thousand Palms Oasis), hereafter referred to as candidate crayfish bioindicators. Candidate bioindicator taxa were from diverse microbial lineages, originating from three kingdoms, 43 orders, and 56 families (Figure 2a, Supplemental Table S4). Each individual bioindicator taxon was found in a minimum of two samples and a minimum of 30 total sequence reads. After conducting the GLOBI search, literature review, and applying qualitative scoring, 60 of these taxa had a weight above 0 but less than 5 (indicating weak to moderate crayfish association), and 9 taxa had a weight of 5 or above, indicating tight crayfish association (Table 2, Supplemental Table S4).

**Figure 2.**
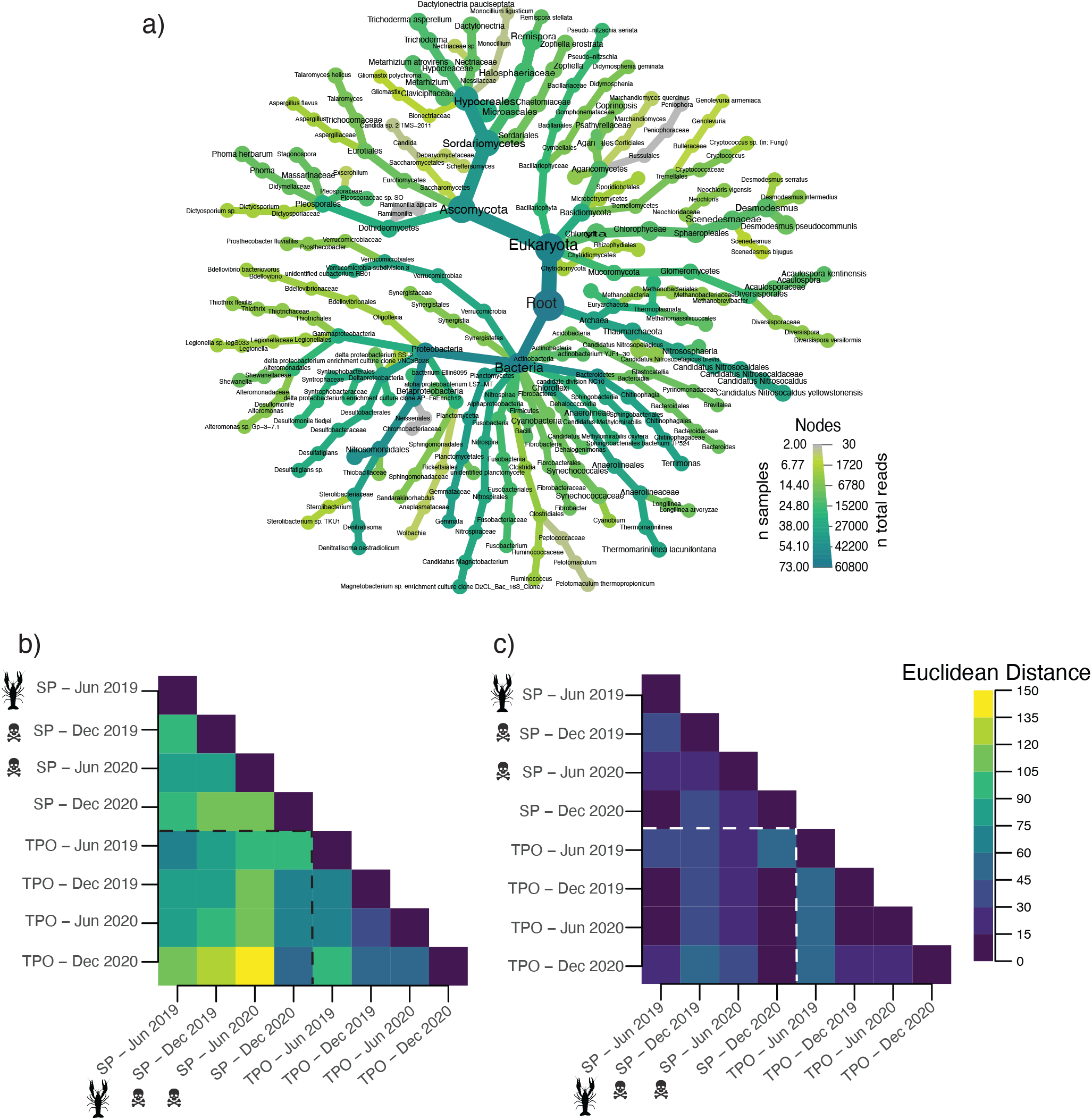
a) Phylogenetic heat tree of 90 crayfish bioindicator species, made using the *Metacoder* R package (Foster et al., 2017). Nodes and branches with higher thickness had higher numbers of sequence reads, darker teal were found in more samples. b,c) Heat maps of frequency of b) 90 bioindicator taxa and c) 90 taxa sampled randomly from 16S and FITS datasets with 20 permutations. Color indicates Euclidean distance in frequency and composition of taxa between sample points, with blue and purple indicating more similarity in composition and green and yellow indicating more differences. Square areas within the dashed lines are comparisons between Simone Pond and Thousand Palms Oasis. Areas outside the dashed lines are comparisons within the same oasis.

**Table 2.** Highly ranked crayfish associates (N=9) ordered by qualitative rank value

| Domain | Phylum | Class | Order | Family | Genus | Species | Rank Value |
| --- | --- | --- | --- | --- | --- | --- | --- |
| Bacteria | Cyanobacteria | Cyanophyceae | Synechococcales | Synechococcaceae | Cyanobium | unknown | 6 |
| Eukaryota | Ascomycota | Eurotiomycetes | Eurotiales | Aspergillaceae | Aspergillus | *Aspergillus flavus* | 6 |
| Eukaryota | Ascomycota | Eurotiomycetes | Eurotiales | Trichocomaceae | Talaromyces | *Talaromyces helicus* | 6 |
| Eukaryota | Ascomycota | Dothideomycetes | Pleosporales | Didymellaceae | Phoma | *Phoma herbarum* | 5 |
| Eukaryota | Ascomycota | Dothideomycetes | Pleosporales | Massarinaceae | Stagonospora | unknown | 5 |
| Eukaryota | Chlorophyta | Chlorophyceae | Sphaeropleales | Scenedesmaceae | Desmodesmus | *Desmodesmus intermedius* | 5 |
| Eukaryota | Chlorophyta | Chlorophyceae | Sphaeropleales | Scenedesmaceae | Desmodesmus | *Desmodesmus pseudocommunis* | 5 |
| Eukaryota | Chlorophyta | Chlorophyceae | Sphaeropleales | Scenedesmaceae | Desmodesmus | *Desmodesmus serratus* | 5 |
| Eukaryota | Chlorophyta | Chlorophyceae | Sphaeropleales | Scenedesmaceae | Scenedesmus | *Scenedesmus bijugus* | 5 |

Heatmap visualizations of Euclidean distance of composition and frequency of 90 candidate bioindicator taxa between oases and throughout time indicate that composition of bioindicator taxa differed between and within oases in both 2019 timepoints and in June 2020 (Figure 2b), indicating strong community turnover of bioindicator taxa during the study period, especially between oases when crayfish were present and when cypermethrin was applied. The same heatmap analysis conducted on 90 random taxa (averaged frequency from 20 permutations) from the total microbial data set (merged 16S and FITS) showed a substantially different pattern than the heatmap made with only bioindicator taxa, with fewer community differences between oases and timepoints (Figure 2c).

#### 3.2.2. SIMPER Analysis of Simone Pond and Thousand Palms Oasis After Crayfish Removal

The SIMPER analyses for the remaining sampling points had fewer taxa return as significant drivers of beta diversity between the two oases, indicating that communities between the two oases may be becoming more similar over time post-crayfish eradication. In the December 2019 samples, taken 5 months into the eradication program, we only detected four taxa that were significantly more likely to occur in Simone Pond as compared to Thousand Palms Oasis. We found no taxa more significantly likely to occur in Simone Pond in June 2020, and one significant taxon in Dec 2020 (Table 3). In December 2019, two of the original bioindicator taxa (crayfish association rank of 0) were significantly more likely to occur in Simone Pond in both June 2019 and Dec 2019 samples (Table 3, bolded text).

**Table 3.**
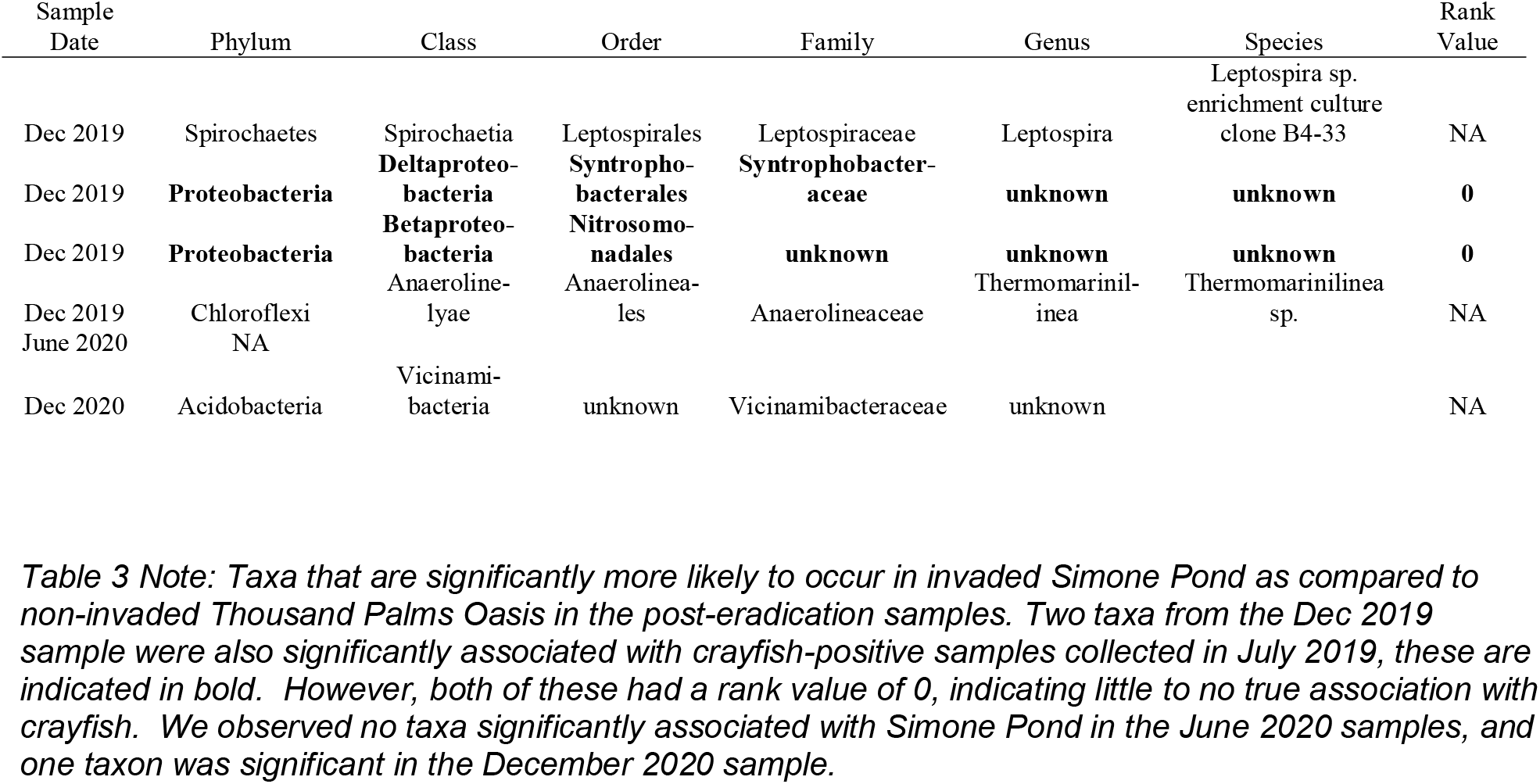
SIMPER Significant Taxa After Crayfish Removal.

#### 3.2.3. Discovery of candidate bioindicator taxa in crayfish carapace sample

We looked for the presence of candidate bioindicator taxa in the microbial metagenome of the crayfish carcass (Section 3.1.3) and found strong overlap (Supplemental Table S3). At the family level, we found 24 overlapping families with the 90 bioindicators, encompassing 32 bioindicator taxa. At the genus level, we found 17 overlapping genera encompassing 18 bioindicator taxa including top ranked bioindicators (rank 5 or 6): *Cyanobium sp, Aspergillus, Talaromyces*, and *Scenedesmus*. *Shewanella*, a taxon that we did not rank highly in our qualitative score (rank 2) but that is documented to make up 20% of the *P. clarkii* crayfish gut microbiome (Shui et al., 2020), was the dominant of the overlapping genera with 20,106 reads and the third highest in overall relative abundance in the carcass metagenome. These bioindicators were found at no such level in the sample periods following crayfish removal (Supplemental Table S1).

### 3.3. Individual Metabarcode Biodiversity Analysis

#### 3.3.1 Metabarcode Sequencing Summary

We generated 47.0 million MiSeq paired reads and 52.6 million NextSeq paired reads. Negative controls including field blanks, extraction blanks, and PCR blanks made up 2 million of those MiSeq reads and 3.7 million of the NextSeq reads (NCBI bioproject, will provide reference after manuscript acceptance). The *16S* marker targeting Prokaryota recovered 2280 taxa in June 2019 samples from MiSeq sequencing and in subsequent sampling windows of December 2019, June 2020, and December 2020, which used the NextSeq for less costly sequencing, we recovered 2979, 3899, and 2614 taxa, respectively. 91% of these taxa were Bacteria, and 5% were Archaea and 4% Eukaryota. In the same sample date order, the *18S* marker targeting Eukaryota recovered 1668, 1213, 1041, and 1049 taxa and all were assigned to Eukaryota. The *FITS* marker targeting Fungi recovered only Eukaryota, and included 636, 605, 967, and 649 taxa. The full *CO1* marker targeting animals and protists used in June 2019 recovered 427 taxa and contained 76% Bacteria, 0.7% Archaea, and 23% Eukaryota. The partial CO1 marker used in subsequent sampling amplified 916, 760, and 513 taxa consisting of over 99% Eukaryota and 0.3% Bacteria. The *PITS* marker targeting Chlorophyta (land plants and green algae) recovered 125, 141, 145, and 100 taxa and all were Eukaryota, of which 66% belonged to Streptophyta and the remainder to Chlorophyta.

#### 3.3.2 Beta Diversity Analyses

For four of the marker datasets (18S - Eukaryotes, FITS - Fungi, 16S - Bacteria and Archaea, and PITS - Plants), Simone Pond and Thousand Palms Oasis had significantly different community composition from each other at each sample time point (PERMANOVA p<0.05, Figure 3a, Supplemental Figure S2a-c, Supplemental Table S5). PERMANOVA analyses for the CO1 marker set indicated that communities were significantly different at the latter three sample periods, but not in June 2019 (Figure S2d), which was due to Proteobacteria swamping, an unintended effect of using the full-length COI primers for the June 2019 dataset.

**Figure 3.**
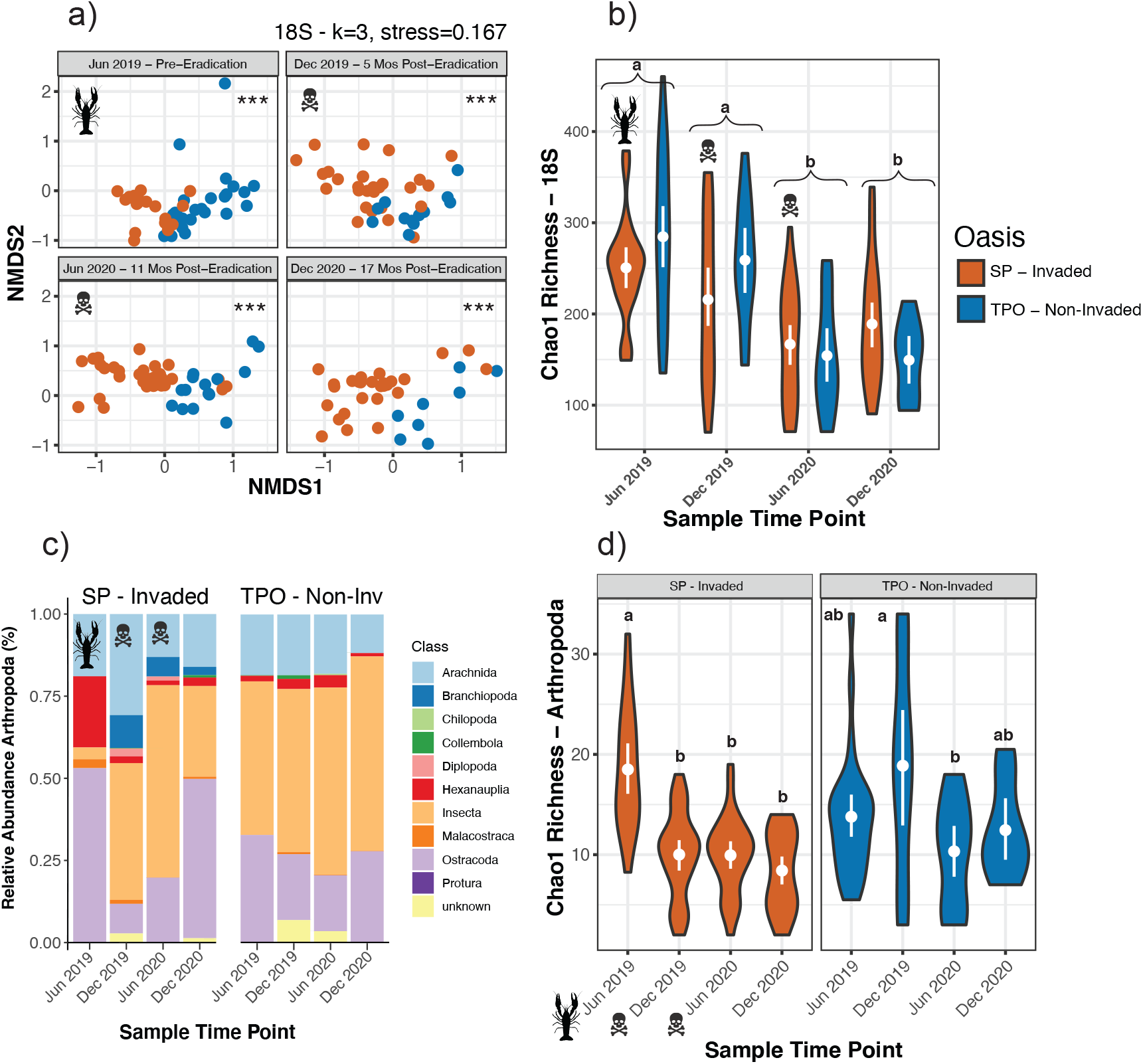
Biodiversity within a, b) the full 18S (Eukaryote) dataset and c, d) Arthropoda only from 18S and CO1 combined datasets. Thousand Palms Oasis (TPO) is represented in blue and Simone Pond (SP) is represented in orange for panels a, b, and d. a) Axes 1 and 2 of NMDS ordination faceted by sample date. b) Violin plot of Chao1 richness at each sample point. Internal white points and bars indicate mean value with 95% confidence interval generated from 1000 bootstraps. Letters indicate significant group differences in species richness by sample date (group a or b). There was no difference in species richness between oases within a timepoint. c) Barplot showing relative sequence read abundance of detected classes of Arthropoda in each sample and oasis. d) Violin plot of Chao1 richness of Arthropoda at each sample point, faceted by oasis. Internal white points and bars indicate mean value with 95% confidence interval generated from 1000 bootstraps. Letters indicate significant group differences in species richness by oasis and sample date (group a or b, within an oasis).

Pairwise PERMANOVA tests also indicated differences in community composition within oases between different sample time periods. However, the time periods that significantly differed within an oasis was variable within and between datasets (Supplemental Table S5). For instance, Simone Pond 18S eukaryotic communities in Jun 2020 and Dec 2020 were significantly different, but in Thousand Palms Oasis they were not. The test for differences in beta dispersion for Simone Pond 18S was significant when comparing Jun 2019 and Dec 2019 samples (F=4.06, p=0.012) indicating that significant differences in these two sampling points shown by PERMANOVA could be due to differences in dispersion. All other 18S community comparisons from Simone Pond and Thousand Palms Oasis were not significantly different in beta dispersion, indicating that PERMANOVA results are accurately reflecting a true difference in community structure between remaining time points. Full PERMANOVA and Beta Dispersion Test results for all marker sets are reported in Supplemental Table S5.

#### 3.3.3 Overall Alpha Diversity Analyses

Alpha diversity estimates were more variable between individual marker datasets, with different taxonomic groups showing different levels of species richness between oases and timepoints. In general, ANOVA with post-hoc Tukey tests indicated that extrapolated species richness (Chao1) at individual time points within each dataset did not significantly differ between Oases (Figure 3b, Supplemental Figure S3). There were two exceptions to these results, where Bacteria and Archaea (16S) in Simone Pond 16S showed significantly lower species richness than Thousand Palms Oasis in Jun 2019 (Supplemental Figure S3b, Supplemental Table S6), and Plants (PITS) in Simone Pond having significantly higher species richness than Thousand Palms Oasis in Dec 2019 (Supplemental Figure S3c).

When species richness of both Oases was considered together, each marker dataset showed significant differences between timepoints, although there was not any time point that was consistently more species-rich. For 18S Eukaryotes, Jun 2019 and Dec 2019 had the highest species richness for both oases (Figure 3b); for 16S, PITS, and CO1, Dec 2019 showed the highest levels of species richness (Supplemental Figure S3b-d); and Jun 2020 showed the highest species richness for FITS (Supplemental Figure S3a).

#### 3.3.4. Arthropod Biodiversity Analyses

We conducted alpha and beta diversity analyses of only Arthropoda taxa by merging 18S Eukaryotes and CO1 Invertebrates and Protists datasets and removing all non-arthropod taxa. We observed a significant effect of sample date on arthropod species richness in Simone Pond (ANOVA, F=13.85, p<0.001). Post-hoc Tukey tests indicate that all samples at time points after crayfish eradication in Simone Pond had lower observed alpha diversity than the Jun 2019 sample pre-eradication (p<0.001, Figure 2d). In Thousand Palms Oasis, arthropod diversity was also significantly affected by sample period (ANOVA, F=3.84, p=0.019), but Tukey tests reveal that arthropod species richness in Thousand Palms Oasis remained more stable between sample points, with only the June 2020 sample significantly lower than the December 2019 sample (p=0.023, Figure 3d). We found no other significant difference in species richness between any sample point in Thousand Palms Oasis (see Supplemental Table S6 for full ANOVA and post-hoc test results).

PERMANOVA analyses indicate significant arthropod community turnover across time in both Simone Pond and Thousand Palms Oasis (Table S5, Figure 3c). In Thousand Palms Oasis, all sample periods had significantly different communities from each other except for June 2020 and December 2020, similar to results for the single marker datasets described above. All sample points had significantly different communities in Simone Pond and higher F values than comparisons between Thousand Palms Oasis, indicating stronger separation of arthropod communities between sample points in Simone Pond than Thousand Palms Oasis.

## 4. Discussion

### 4.1 Candidate bioindicators of crayfish presence are valuable to monitor over time

Our analyses show individual taxa as significant drivers of community difference between crayfish positive and negative samples, which allows for flagging of potential bioindicator species at different times and between invaded and non-invaded sites. We found that several microbial taxa identified through eDNA metabarcoding are reliable bioindicators for invasive crayfish. SIMPER analyses of the two oases and experimental tanks identified 90 taxa that were significantly more likely to occur when crayfish were present. After conducting literature review and qualitative scoring on these taxa, we found 9 microbial taxa that are highly linked to crayfish. We propose these 9 taxa as candidate crayfish bioindicators that can be used as potential warning signs of invasive crayfish presence in our study site as well as other systems where crayfish are suspected or known to be present. We found that many of these taxa are directly linked to crayfish as a specialized host, and others that are likely associated with crayfish due to shared habitat requirements. Our highest ranked candidate bioindicators, with rank values of 6 (*Cynobium* sp., *Aspergillus flavus*, and *Talaromyces helicus)* were previously identified as key associated taxa in invasive *P. clarkii* or *C. chelax* in Italy and Australia (Dörr et al., 2012; Foysal et al., 2019; Garzoli et al., 2014). In our study, out of 1003 total eDNA samples from across the state of California (Meyer et al. 2021), *Cynobium sp*. and *Talaromyces helicus* were only found in samples in the Coachella Valley, suggesting that they may thrive in desert environments where crayfish are present. *Aspergillius flavus, Phoma herbarum* and several of the mid-ranked biodindicators in our study have been previously classified as fungal plant pathogens (Dörr et al., 2012; e.g. Stagnospora sp., Liu et al., 2012; Ramimonilia apicalis, Ricks, 2019). Garzoli et al. (2014) suggested that these and other fungal species are common in the crayfish gut to help break down plant detritus; a primary source of food for *P. clarkii*, and thus are likely to be tightly associated with *P. clarkii*. Overall, 4 out of 9 candidate bioindicator taxa were fungal species in the Phylum Ascomycota. This finding is consistent with the hypothesis that invasive crayfish may act as vectors for fungal plant pathogens (Dörr et al. 2012; Garzoli et al. 2014). The remaining four taxa in the 9 candidate bioindicators are all green algae in the family Scenedesmaceae. Congeners of these taxa have been previously found to co-occur with crayfish, and have been shown to be sensitive to crayfish pesticides (Buřič et al., 2013; Sarıkaya et al., 2011), suggesting that they may constitute important habitat for *P. clarkii*. We note that invasive tilapia were also present in our study system, potentially inflating the number of taxa significantly associated with crayfish in the pre-eradication samples. However, given that we included experimental tanks containing only tilapia as non-crayfish samples, we are confident that any tilapia-specific taxa were filtered out of our final data set of crayfish-associated taxa. We suggest that the microbial taxa reported here constitute a proxy indicator community for crayfish presence, which we propose as a valuable addition to a multi-pronged detection approach along with visual, trapping, and/or direct molecular surveys, as per Tregulier et al. (2014).

It is unclear why multiple methods of direct molecular detection of crayfish failed in our study. It is possible that the timing of our sample did not coincide with a period when crayfish shed the most DNA (e.g. during molting or gravidity, Dunn et al. 2017), and so the levels of DNA in both the oasis and tank samples were not high enough to be detected at high confidence. Additionally, the published primer studies noting successful detection of *P. clarkii* from environmental samples used crayfish samples in Europe, so it is also possible that there are mutations in primer binding sites between these geographically disparate populations. Genome sequencing of multiple *P. clarkii* populations around the globe would be useful to understand if sequence differences are contributing to unstable molecular detection, but this was not within the scope of our study. Regardless, the variability of detection success of *P. clarkii* both within and between studies suggest that a multi-pronged approach including some combination of visual surveys, trapping, and molecular detection, including direct (qPCR, targeted PCR) and/or indirect (metabarcoding, microbial proxies) is necessary for the management of this species. We suggest that for managers using molecular surveys, microbial indicators can be tracked during the same analysis. If either crayfish or microbial indicators are found in the sequencing data, managers can be on alert for potential invasion or crayfish refugia.

### 4.2 Community composition but not species richness varies between oases through time

Biodiversity analyses of five metabarcode loci (18S, eukaryotes; FITS, fungi; 16S, Bacteria and Archaea; CO1, animals; and PITS, plants) showed that community composition between oases in all taxa was significantly different at each sample period (with the exception of CO1 in Jun 2019 due to proteobacteria contamination). Communities continued to show significant differences even in the final sample period (December 2020), which was 17 months post-eradication and 5 months post-pesticide application in Simone Pond. This result is somewhat surprising as both oases are fed from the same water source, are located less than 0.5 km apart, and are characterized by the same broad vegetation type (California Fan Palm, *Washingtonia filifera)*. Due to the legacy of invasive crayfish and tilapia in Simone Pond, we expected that community composition would be different between Simone Pond and Thousand Palms Oasis in the pre-eradication sample but that the two sites would become more similar over time. However, our results indicate that Simone Pond may still be recovering from invasive presence and eradication. Indeed, it is possible that complete ecological recovery in Simone Pond may never occur to the extent that communities between Thousand Palms Oasis and Simone Pond become similar (Prior et al., 2018). It is also possible that Thousand Palms Oasis and Simone Pond support inherently different biological communities, currently and historically. Thousand Palms Oasis has a more closed canopy and shallower depth than Simone Pond, which is why only Simone Pond and not Thousand Palms Oasis is targeted as a re-introduction site for the desert pupfish (Marsh and Sada, 1993; Varela-Romero et al., 2002).

While community composition differed between oases, species richness remained relatively consistent between them. Different datasets showed different peaks of species richness at different time points, but richness rarely differed between oases at a single time point. Again, we were surprised by this result as we expected that invasive presence and eradication would differentially affect overall species richness between invaded and non-invaded oases. This was only true when analyzing the species richness of Arthropoda over time. Other studies have similarly shown that invasive species may not decrease overall species richness, especially when considering native and non-native species (Fork, 2010; Forys and Allen, 2002; Matthews et al., 2009). Tracking both native and overall community composition and richness between oases, along with invasive species bioindicators, will continue to be of interest as the next phases of restoration at the preserve continue (Short, 2017).

### 4.3 Cypermethrin negatively impacted arthropod biodiversity in Simone Pond

Although the pesticide cypermethrin was deemed necessary to eradicate crayfish from Simone Pond, managers were concerned about the potential for off-target effects of the pesticides on other arthropod taxa. Our analyses of taxa in Arthropoda in CO1 and 18S datasets reveal that all three sample periods post-eradication had significantly lower arthropod species richness than the pre-eradication sample in Simone Pond. For example, taxa in the classes Malacostraca (which also include crayfish), and Hexanuplia (containing mainly Copepods) were found in the June 2019 sample in Simone Pond, but were either eliminated or found in lower sequence read abundance in subsequent samples. The final sample in December 2020, where pesticide had not been applied for 5 months, had levels of arthropod species richness that were similar to the earlier post-eradication samples. Analysis of Thousand Palms Oasis did not show the same change in arthropod species richness through time: post-eradication samples had similar or higher levels of arthropod species richness compared to December 2019. As Thousand Palms Oasis was effectively a control sample with no pesticide applied, these results suggest that, while the cypermethrin negatively impacted arthropod species richness in Simone Pond, it did not affect arthropod communities in the other oases in the preserve. Previous studies have highlighted the persistent toxicity of pyrethroid pesticides in sediments on benthic invertebrates (Li et al., 2017; Zhang et al., 2018). Further monitoring of sites treated with cypermethrin for the purposes of crayfish eradication will be necessary to understand the long-term effects of cypermethrin treatment on both aquatic and terrestrial arthropod communities.

## 5. Conclusions and Conservation Applications

We used multi-locus metabarcoding of environmental DNA in an aquatic ecosystem undergoing conservation management to simultaneously identify microbial bioindicators for an invasive crayfish and track biodiversity changes. We were unable to reliably detect crayfish DNA in environmental samples using either metabarcoding or qPCR, which is consistent with previous studies of crayfish (Treguier et al. 2014) and other crustaceans (e.g. Crane et al. 2021). However, our community-level analyses of bacterial and fungal markers in habitats where crayfish were known to be present (in tanks with crayfish and the invaded oasis pre-eradication) revealed a community of microbial taxa that are strongly associated with crayfish presence. Analyses of post-eradication environmental samples from the invaded pond revealed few of these bioindicator taxa, suggesting that crayfish are no longer present, consistent with the traditional survey data. Our biodiversity analyses of five metabarcode datasets also revealed that biological communities differed between oases through time, although species richness estimates tended to be similar between oases at individual time points. Finally, we found that arthropod diversity declined after crayfish pesticide treatment but did not decline in the oasis that was not treated with pesticide, suggesting that cypermethrin use may have detrimental off-target effects to other species. Overall, we recommend conducting ongoing monitoring using multiple methods including eDNA for native and invasive species as a powerful approach to track biodiversity and invasive status throughout a conservation project. As managers of invaded systems look for solutions for restoration and monitoring, the use of metabarcoding can be an invaluable tool to track cryptic species with multiple approaches and monitor biodiversity.

## Acknowledgments

The authors wish to acknowledge Deborah Rogers, Center for Natural Lands Management (CNLM) Director of Conservation Science and Stewardship, for providing invaluable feedback on manuscript drafts. We thank C. Orland, H. Bregoff, C.W. Schaal, A. Monzón, E. De Garcas, R. Henderson, and Y. Xiao for assistance with environmental DNA sampling. We finally wish to thank the members of the CALeDNA team and eDNA working group in the Paleogenomics Lab at the University of California Santa Cruz for overall support, as well as continual feedback on data analysis and visualization throughout the project. This work was funded by the HHMI Professors Program grant no. GT10483 as well as the State of California Proposition 1 and USFWS Strategic Habitat Conservation Grants.

## References

Alcorlo, P., Geiger, W., Otero, M., 2004. Feeding Preferences and Food Selection of the Red Swamp Crayfish, Procambarus clarkii, in Habitats Differing in Food Item Diversity. Crustaceana 77, 435–453.

Amaral-Zettler, L.A., McCliment, E.A., Ducklow, H.W., Huse, S.M., 2009. A Method for Studying Protistan Diversity Using Massively Parallel Sequencing of V9 Hypervariable Regions of Small-Subunit Ribosomal RNA Genes. PLOS ONE 4, e6372. https://doi.org/10.1371/journal.pone.0006372

Anderson, M.J., Walsh, D.C.I., 2013. PERMANOVA, ANOSIM, and the Mantel test in the face of heterogeneous dispersions: What null hypothesis are you testing? Ecological Monographs 83, 557–574. https://doi.org/10.1111/j.2041-210X.2011.00127.x

Astudillo-García, C., Hermans, S.M., Stevenson, B., Buckley, H.L., Lear, G., 2019. Microbial assemblages and bioindicators as proxies for ecosystem health status: potential and limitations. Appl Microbiol Biotechnol 103, 6407–6421. https://doi.org/10.1007/s00253-019-09963-0

Ballare, K.M., Neff, J.L., Ruppel, R., Jha, S., 2019. Multi-scalar drivers of biodiversity: local management mediates wild bee community response to regional urbanization. Ecological Applications 29, e01869.

Barnes, M.A., Turner, C.R., Jerde, C.L., Renshaw, M.A., Chadderton, W.L., Lodge, D.M., 2014. Environmental Conditions Influence eDNA Persistence in Aquatic Systems. Environ. Sci. Technol. 48, 1819–1827. https://doi.org/10.1021/es404734p

Bucciarelli, G.M., Suh, D., Lamb, A.D., Roberts, D., Sharpton, D., Shaffer, H.B., Fisher, R.N., Kats, L.B., 2019. Assessing effects of non-native crayfish on mosquito survival. Conservation Biology 33, 122–131. https://doi.org/10.1111/cobi.13198

Buřič, M., Kouba, A., Máchová, J., Mahovská, I., Kozák, P., 2013. Toxicity of the organophosphate pesticide diazinon to crayfish of differing age. Int. J. Environ. Sci. Technol. 10, 607–610. https://doi.org/10.1007/s13762-013-0185-4

Bushnell, B., 2014. BBTools. BBMap short read aligner, and other bioinformatic tools. Available online at: https://sourceforge.net/projects/bbmap.

Cai, P., Huang, Q., Zhang, X., Chen, H., 2006. Adsorption of DNA on clay minerals and various colloidal particles from an Alfisol. Soil Biology and Biochemistry 38, 471–476. https://doi.org/10.1016/j.soilbio.2005.05.019

Cai, W., Ma, Z., Yang, C., Wang, L., Wang, W., Zhao, G., Geng, Y., Yu, D.W., 2017. Using eDNA to detect the distribution and density of invasive crayfish in the Honghe-Hani rice terrace World Heritage site. PLOS ONE 12, e0177724. https://doi.org/10.1371/journal.pone.0177724

Callahan, B.J., McMurdie, P.J., Rosen, M.J., Han, A.W., Johnson, A.J.A., Holmes, S.P., 2016. DADA2: High-resolution sample inference from Illumina amplicon data. Nature methods 13, 581–583.

Caporaso, J.G., Lauber, C.L., Walters, W.A., Berg-Lyons, D., Huntley, J., Fierer, N., Owens, S. M., Betley, J., Fraser, L., Bauer, M., Gormley, N., Gilbert, J.A., Smith, G., Knight, R., 2012. Ultra-high-throughput microbial community analysis on the Illumina HiSeq and MiSeq platforms. ISME J 6, 1621–1624. https://doi.org/10.1038/ismej.2012.8

Chen, X., Fan, L., Qiu, L., Dong, X., Wang, Q., Hu, G., Meng, S., Li, D., Chen, J., 2021. Metagenomics Analysis Reveals Compositional and Functional Differences in the Gut Microbiota of Red Swamp Crayfish, Procambarus clarkii, Grown on Two Different Culture Environments. Frontiers in Microbiology 12.

Chucholl, F., Fiolka, F., Segelbacher, G., Epp, L.S., 2021. eDNA Detection of Native and Invasive Crayfish Species Allows for Year-Round Monitoring and Large-Scale Screening of Lotic Systems. Frontiers in Environmental Science 9.

Clarke, K.R., 1993. Non-parametric multivariate analyses of changes in community structure. Australian Journal of Ecology 18, 117–143.

Crane, L.C., Goldstein, J.S., Thomas, D.W., Rexroth, K.S., Watts, A.W., 2021. Effects of life stage on eDNA detection of the invasive European green crab (Carcinus maenas) in estuarine systems. Ecological Indicators 124, 107412. https://doi.org/10.1016/j.ecolind.2021.107412

Curd, E.E., Gold, Z., Kandlikar, G.S., Gomer, J., Ogden, M., O’Connell, T., Pipes, L., Schweizer, T. M., Rabichow, L., Lin, M., Shi, B., Barber, P.H., Kraft, N., Wayne, R., Meyer, R.S., 2019. Anacapa Toolkit: An environmental DNA toolkit for processing multilocus metabarcode datasets. Methods in Ecology and Evolution 10, 1469–1475.

Curtis, A.N., Larson, E.R., 2020. No evidence that crayfish carcasses produce detectable environmental DNA (eDNA) in a stream enclosure experiment. PeerJ 8, e9333. https://doi.org/10.7717/peerj.9333

Davis, N.M., Proctor, D.M., Holmes, S.P., Relman, D.A., Callahan, B.J., 2018. Simple statistical identification and removal of contaminant sequences in marker-gene and metagenomics data. Microbiome 6, 1–14.

Dörr, A.J.M., Rodolfi, M., Elia, A.C., Scalici, M., Garzoli, L., Picco, A.M., 2012. Mycoflora on the cuticle of the invasive crayfish Procambarus clarkii. Fundamental and Applied Limnology 180, 77–84.

Dougherty, M.M., Larson, E.R., Renshaw, M.A., Gantz, C.A., Egan, S.P., Erickson, D.M., Lodge, D.M., 2016. Environmental DNA (eDNA) detects the invasive rusty crayfish Orconectes rusticus at low abundances. Journal of Applied Ecology 53, 722–732. https://doi.org/10.1111/1365-2664.12621

Dragičević, P., Bielen, A., Petrić, I., Hudina, S., 2021. Microbial pathogens of freshwater crayfish: A critical review and systematization of the existing data with directions for future research. Journal of Fish Diseases 44, 221–247. https://doi.org/10.1111/jfd.13314

Dunn, N., Priestley, V., Herraiz, A., Arnold, R., Savolainen, V., 2017. Behavior and season affect crayfish detection and density inference using environmental DNA. Ecology and Evolution 7, 7777–7785. https://doi.org/10.1002/ece3.3316

Edwards, R., Millburn, P., Hutson, D.H., 1986. Comparative toxicity of cis-cypermethrin in rainbow trout, frog, mouse, and quail. Toxicology and Applied Pharmacology 84, 512–522. https://doi.org/10.1016/0041-008X(86)90256-5

Ficetola, G.F., Manenti, R., Taberlet, P., 2019. Environmental DNA and metabarcoding for the study of amphibians and reptiles: species distribution, the microbiome, and much more. Amphibia-Reptilia 40, 129–148. https://doi.org/10.1163/15685381-20191194

Ficetola, G.F., Miaud, C., Pompanon, F., Taberlet, P., 2008. Species detection using environmental DNA from water samples. Biology Letters 4, 423–425. https://doi.org/10.1098/rsbl.2008.0118

Fork, S.K., 2010. Arthropod Assemblages on Native and Nonnative Plant Species of a Coastal Reserve in California. Environ Entomol 39, 753–762. https://doi.org/10.1603/EN09185

Forys, E.A., Allen, C.R., 2002. Functional Group Change within and across Scales following Invasions and Extinctions in the Everglades Ecosystem. Ecosystems 5, 339–347. https://doi.org/10.1007/s10021-001-0078-0

Foster, Z.S.L., Sharpton, T.J., Grünwald, N.J., 2017. Metacoder: An R package for visualization and manipulation of community taxonomic diversity data. PLOS Computational Biology 13, e1005404. https://doi.org/10.1371/journal.pcbi.1005404

Foysal, M.J., Nguyen, T.T.T., Chaklader, M.R., Siddik, M.A.B., Tay, C.-Y., Fotedar, R., Gupta, S.K., 2019. Marked variations in gut microbiota and some innate immune responses of fresh water crayfish, marron (Cherax cainii, Austin 2002) fed dietary supplementation of Clostridium butyricum. PeerJ 7, e7553. https://doi.org/10.7717/peerj.7553

Gao, X., Lin, H., Revanna, K., Dong, Q., 2017. A Bayesian taxonomic classification method for 16S rRNA gene sequences with improved species-level accuracy. BMC bioinformatics 18, 1–10.

Garzoli, L., Paganelli, D., Rodolfi, M., Savini, D., Moretto, M., Occhipinti, A., Picco, A., 2014. First evidence of microfungal “extra oomph” in the invasive red swamp crayfish Procambarus clarki. Aquatic Invasions 9, 47–58. https://doi.org/10.3391/ai.2014.9.1.04

Geerts, A.N., Boets, P., Van den Heede, S., Goethals, P., Van der heyden, C., 2018. A search for standardized protocols to detect alien invasive crayfish based on environmental DNA (eDNA): A lab and field evaluation. Ecological Indicators 84, 564–572. https://doi.org/10.1016/j.ecolind.2017.08.068

Geiger, W., Alcorlo, P., Baltanás, A., Montes, C., 2005. Impact of an introduced Crustacean on the trophic webs of Mediterranean wetlands. Biological Invasions 7, 49–73. https://doi.org/10.1007/s10530-004-9635-8

Gordon, A., Hannon, G., 2010. Fastx-toolkit. FASTQ/A short-reads preprocessing tools (unpublished) http://hannonlab.cshl.edu/fastx_toolkit 5.

Gu, W., Song, J., Cao, Y., Sun, Q., Yao, H., Wu, Q., Chao, J., Zhou, J., Xue, W., Duan, J., 2013. Application of the ITS2 Region for Barcoding Medicinal Plants of Selaginellaceae in Pteridophyta. PLOS ONE 8, e67818. https://doi.org/10.1371/journal.pone.0067818

Halstead, B.J., Wood, D.A., Bowen, L., Waters, S.C., Vandergast, A.G., Ersan, J.S., Skalos, S.M., Casazza, M.L., 2017. An evaluation of the efficacy of using environmental DNA (eDNA) to detect giant gartersnakes (Thamnophis gigas) (No. 2017–1123), Open-File Report. U.S. Geological Survey. https://doi.org/10.3133/ofr20171123

Harper, K.J., Anucha, P., Turnbull, J.F., Bean, C.W., Leaver, M.J., 2018. Searching for a signal: environmental DNA (eDNA) for the detection of invasive signal crayfish, Pacifasticus leniusculus (Dana, 1852). Management of Biological Invasions 9, 137–148. https://doi.org/10.3391/mbi.2018.9.2.07

Hebert, P.D.N., Cywinska, A., Ball, S.L., deWaard, J.R., 2003. Biological Identifications through DNA Barcodes. Proceedings: Biological Sciences 270, 313–321.

Ikeda, K., Doi, H., Tanaka, K., Kawai, T., Negishi, J.N., 2016. Using environmental DNA to detect an endangered crayfish Cambaroides japonicus in streams. Conservation Genet Resour 8, 231–234. https://doi.org/10.1007/s12686-016-0541-z

Jarman, S.N., Berry, O., Bunce, M., 2018. The value of environmental DNA biobanking for long-term biomonitoring. Nature Ecology & Evolution 2, 1192–1193. https://doi.org/10.1038/s41559-018-0614-3

Kandlikar, G.S., Gold, Z.J., Cowen, M.C., Meyer, R.S., Freise, A.C., Kraft, N.J., Moberg-Parker, J., Sprague, J., Kushner, D.J., Curd, E.E., 2018. ranacapa: An R package and Shiny web app to explore environmental DNA data with exploratory statistics and interactive visualizations. F1000Research 7.

Khan, A., Faridi, H.A.M., Ali, M., Khan, M.Z., Siddique, M., Hussain, I., Ahmad, M., 2009. Effects of cypermethrin on some clinico-hemato-biochemical and pathological parameters in male dwarf goats (Capra hircus). Experimental and Toxicologic Pathology 61, 151–160. https://doi.org/10.1016/j.etp.2008.07.001

Knudsen, S.W., Agersnap, S., Møller, P.R., Andersen, J.H., 2019. Development of species-specific eDNA-based test systems for monitoring of freshwater crayfish. NIVA-rapport.

Langmead, B., Salzberg, S.L., 2012. Fast gapped-read alignment with Bowtie 2. Nature methods 9, 357–359.

Leray, M., Yang, J.Y., Meyer, C.P., Mills, S.C., Agudelo, N., Ranwez, V., Boehm, J.T., Machida, R.J., 2013. A new versatile primer set targeting a short fragment of the mitochondrial COI region for metabarcoding metazoan diversity: application for characterizing coral reef fish gut contents. Frontiers in Zoology 10, 34. https://doi.org/10.1186/1742-9994-10-34

Leslie, P.H., Davis, D.H.S., 1939. An Attempt to Determine the Absolute Number of Rats on a Given Area. Journal of Animal Ecology 8, 94–113. https://doi.org/10.2307/1255

Li, H., Cheng, F., Wei, Y., Lydy, M.J., You, J., 2017. Global occurrence of pyrethroid insecticides in sediment and the associated toxicological effects on benthic invertebrates: An overview. Journal of Hazardous Materials 324, 258–271. https://doi.org/10.1016/j.jhazmat.2016.10.056

Li, H., Durbin, R., 2009. Fast and accurate short read alignment with Burrows–Wheeler transform. bioinformatics 25, 1754–1760.

Liu, Z., Zhang, Z., Faris, J.D., Oliver, R.P., Syme, R., McDonald, M.C., McDonald, B.A., Solomon, P.S., Lu, S., Shelver, W.L., Xu, S., Friesen, T.L., 2012. The Cysteine Rich Necrotrophic Effector SnTox1 Produced by Stagonospora nodorum Triggers Susceptibility of Wheat Lines Harboring Snn1. PLOS Pathogens 8, e1002467. https://doi.org/10.1371/journal.ppat.1002467

Marsh, P.C., Sada, D.W., 1993. Draft Desert Pupfish RP Amendment.pdf. U.S. Fish and Wildlife Service.

Martin, B.A., Saiki, M.K., 2005. Relation of desert pupfish abundance to selected environmental variables in natural and manmade habitats in the Salton Sea basin. Environ Biol Fish 73, 97–107. https://doi.org/10.1007/s10641-004-5569-3

Martin, M., 2011. Cutadapt removes adapter sequences from high-throughput sequencing reads. EMBnet.journal 17, 10–12. https://doi.org/10.14806/ej.17.1.200

Matthews, J.W., Peralta, A.L., Soni, A., Baldwin, P., Kent, A.D., Endress, A.G., 2009. Local and landscape correlates of non-native species invasion in restored wetlands. Ecography 32, 1031–1039. https://doi.org/10.1111/j.1600-0587.2009.05863.x

Mauvisseau, Q., Coignet, A., Delaunay, C., Pinet, F., Bouchon, D., Souty-Grosset, C., 2018. Environmental DNA as an efficient tool for detecting invasive crayfishes in freshwater ponds. Hydrobiologia 805, 163–175. https://doi.org/10.1007/s10750-017-3288-y

McMurdie, P.J., Holmes, S., 2012. Phyloseq: a bioconductor package for handling and analysis of high-throughput phylogenetic sequence data, in: Biocomputing 2012. World Scientific, pp. 235–246.

Meyer, F., Paarmann, D., D’Souza, M., Olson, R., Glass, E., Kubal, M., Paczian, T., Rodriguez, A., Stevens, R., Wilke, A., Wilkening, J., Edwards, R., 2008. The metagenomics RAST server – a public resource for the automatic phylogenetic and functional analysis of metagenomes. BMC Bioinformatics 9, 386. https://doi.org/10.1186/1471-2105-9-386

Meyer, R.S., Ramos, M.M., Lin, M., Schweizer, T.M., Gold, Z., Ramos, D.R., Shirazi, S., Kandlikar, G., Kwan, W.-Y., Curd, E.E., Freise, A., Parker, J.M., Sexton, J.P., Wetzer, R., Pentcheff, N.D., Wall, A.R., Pipes, L., Garcia-Vedrenne, A., Mejia, M.P., Moore, T., Orland, C., Ballare, K.M., Worth, A., Beraut, E., Aronson, E., Nielson, R., Lewin, H.A., Barber, P.H., Wall, J., Kraft, N., Shapiro, B., Wayne, R.K., 2021. The CALeDNA program: Citizen scientists and researchers inventory California’s biodiversity. California Agriculture 75. https://doi.org/10.3733/ca.2021a0001

Morolli, C., Quaglio, F., Della Rocca, G., Malvisi, J., Di Salvo, A., 2006. EVALUATION OF THE TOXICITY OF SYNTHETIC PYRETHROIDS TO RED SWAMP CRAYFISH (PROCAMBARUS CLARKII, GIRARD 1852) AND COMMON CARP (CYPRINUS CARPIO, L. 1758). Bull. Fr. Pêche Piscic. 1381–1394. https://doi.org/10.1051/kmae:2006042

Nunes, A.L., Hoffman, A.C., Zengeya, T.A., Measey, G.J., Weyl, O.L., 2017. Red swamp crayfish, Procambarus clarkii, found in South Africa 22 years after attempted eradication. Aquatic Conservation: Marine and Freshwater Ecosystems 27, 1334–1340. https://doi.org/10.1002/aqc.2741

Oficialdegui, F.J., Sánchez, M.I., Clavero, M., 2020. One century away from home: how the red swamp crayfish took over the world. Rev Fish Biol Fisheries 30, 121–135. https://doi.org/10.1007/s11160-020-09594-z

Oksanen, J., Blanchet, F.G., Kindt, R., Legendre, P., Minchin, P.R., O’hara, R., Simpson, G.L., Solymos, P., Stevens, M.H.H., Wagner, H., 2015. Vegan: community ecology package. 2015. R package version 2.

Poelen, J.H., Simons, J.D., Mungall, C.J., 2014. Global biotic interactions: An open infrastructure to share and analyze species-interaction datasets. Ecological Informatics 24, 148–159. https://doi.org/10.1016/j.ecoinf.2014.08.005

Prior, K.M., Adams, D.C., Klepzig, K.D., Hulcr, J., 2018. When does invasive species removal lead to ecological recovery? Implications for management success. Biol Invasions 20, 267–283. https://doi.org/10.1007/s10530-017-1542-x

Pritchard, E.G., Chadwick, D.D.A., Patmore, I.R., Chadwick, M.A., Bradley, P., Sayer, C.D., Axmacher, J.C., 2021. The ‘Pritchard Trap’: A novel quantitative survey method for crayfish. Ecological Solutions and Evidence 2, e12070. https://doi.org/10.1002/2688-8319.12070

Ratsch, R., Kingsbury, B.A., Jordan, M.A., 2020. Exploration of Environmental DNA (eDNA) to Detect Kirtland’s Snake (Clonophis kirtlandii). Animals 10, 1057. https://doi.org/10.3390/ani10061057

Ricks, N., 2019. A Metagenomic Approach to Understand Stand Failure in Bromus tectorum. Theses and Dissertations.

Russell, S.L., Pepper-Tunick, E., Svedberg, J., Byrne, A., Castillo, J.R., Vollmers, C., Beinart, R.A., Corbett-Detig, R., 2020. Horizontal transmission and recombination maintain forever young bacterial symbiont genomes. PLOS Genetics 16, e1008935. https://doi.org/10.1371/journal.pgen.1008935

Sandodden, R., Johnsen, S.I., 2010. Eradication of introduced signal crayfish Pasifastacus leniusculus using the pharmaceutical BETAMAX VET.®. AI 5, 75–81. https://doi.org/10.3391/ai.2010.5.1.9

Sarıkaya, R., Sepici-Dinçel, A., Ça□lan Karasu Benli, A., Selvi, M., Erkoç, F., 2011. The acute toxicity of fenitrothion on narrow-clawed crayfish (Astacus leptodactylus Eschscholtz, 1823) in association with biomarkers of lipid peroxidation. Journal of Biochemical and Molecular Toxicology 25, 169–174. https://doi.org/10.1002/jbt.20373

Sepulveda, A.J., Nelson, N.M., Jerde, C.L., Luikart, G., 2020. Are Environmental DNA Methods Ready for Aquatic Invasive Species Management? Trends in Ecology & Evolution 35, 668–678. https://doi.org/10.1016/j.tree.2020.03.011

Shafer, T.J., Meyer, D.A., Crofton, K.M., 2005. Developmental neurotoxicity of pyrethroid insecticides: critical review and future research needs. Environ Health Perspect 113, 123–136. https://doi.org/10.1289/ehp.7254

Short, G., 2021. Desert Pupfish Refuge Habitat Restoration Project. Center for Natural Lands Management.

Short, G., 2017. Ten year management plan for the Thousand Palms Oasis and McCallum Grove Preserve (S004 & S050) and Whitewater Mitigation Area (S049) Thousand Palms, CA. Center for Natural Lands Management.

Shui, Y., Guan, Z.-B., Liu, G.-F., Fan, L.-M., 2020. Gut microbiota of red swamp crayfish Procambarus clarkii in integrated crayfish-rice cultivation model. AMB Express 10, 5. https://doi.org/10.1186/s13568-019-0944-9

Skelton, J., Geyer, K.M., Lennon, J.T., Creed, R.P., Brown, B.L., 2017. Multi-scale ecological filters shape the crayfish microbiome. Symbiosis 72, 159–170. https://doi.org/10.1007/s13199-016-0469-9

Soares, M.P., Oliveira, N., Rebelo, D., Marcondes, S.F., Fernandes, C.E., Domingues, I., Soares, A., Hayd, L., 2019. Cypermethrin-based formulation Barrage® induces histological changes in gills of the Pantanal endemic shrimp Macrobrachium pantanalense. Environmental Toxicology and Pharmacology 67, 66–72. https://doi.org/10.1016/j.etap.2019.01.014

Souty-Grosset, C., Anastácio, P.M., Aquiloni, L., Banha, F., Choquer, J., Chucholl, C., Tricarico, E., 2016. The red swamp crayfish Procambarus clarkii in Europe: Impacts on aquatic ecosystems and human well-being. Limnologica 58, 78–93. https://doi.org/10.1016/j.limno.2016.03.003

Suarez-Menendez, M., Planes, S., Garcia-Vazquez, E., Ardura, A., 2020. Early Alert of Biological Risk in a Coastal Lagoon Through eDNA Metabarcoding. Frontiers in Ecology and Evolution 8.

Thalinger, B., Deiner, K., Harper, L.R., Rees, H.C., Blackman, R.C., Sint, D., Traugott, M., Goldberg, C.S., Bruce, K., 2021. A validation scale to determine the readiness of environmental DNA assays for routine species monitoring. Environmental DNA 3, 823–836. https://doi.org/10.1002/edn3.189

Tréguier, A., Paillisson, J.-M., Dejean, T., Valentini, A., Schlaepfer, M.A., Roussel, J.-M., 2014. Environmental DNA surveillance for invertebrate species: advantages and technical limitations to detect invasive crayfish Procambarus clarkii in freshwater ponds. Journal of Applied Ecology 51, 871–879. https://doi.org/10.1111/1365-2664.12262

Troth, C.R., Burian, A., Mauvisseau, Q., Bulling, M., Nightingale, J., Mauvisseau, C., Sweet, M.J., 2020. Development and application of eDNA-based tools for the conservation of white-clawed crayfish. Science of The Total Environment 748, 141394. https://doi.org/10.1016/j.scitotenv.2020.141394

Turner, C.R., Uy, K.L., Everhart, R.C., 2015. Fish environmental DNA is more concentrated in aquatic sediments than surface water. Biological Conservation, Special Issue: Environmental DNA: A powerful new tool for biological conservation 183, 93–102. https://doi.org/10.1016/j.biocon.2014.11.017

Varela-Romero, A., Ruiz-Campos, G., Yépiz-Velázquez, L.M., Alaníz-García, J., 2002. Distribution, habitat and conservation status of desert pupfish (Cyprinodon macularius) in the Lower Colorado River Basin, Mexico. Reviews in Fish Biology and Fisheries 12, 157–165. https://doi.org/10.1023/A:1025006920052

Warton, D.I., Wright, S.T., Wang, Y., 2012. Distance-based multivariate analyses confound location and dispersion effects. Methods in Ecology and Evolution 3, 89–101. https://doi.org/10.18637/jss.v032.i10

Wei, N., Nakajima, F., Tobino, T., 2018. A Microcosm Study of Surface Sediment Environmental DNA: Decay Observation, Abundance Estimation, and Fragment Length Comparison. Environ. Sci. Technol. 52, 12428–12435. https://doi.org/10.1021/acs.est.8b04956

White, T., Bruns, T., Lee, S., Taylor, J., 1990. Amplification and direct sequencing of fungal ribosomal RNA genes for phylogenetics, in: PCR Protocols: A Guide to Methods and Applications. Academic Press, New York, pp. 315–322.

Wickham, H., Chang, W., Wickham, M.H., 2016. Package ‘ggplot2.’ Create elegant data visualisations using the grammar of graphics. Version 2, 1–189.

Wittwer, C., Stoll, S., Strand, D., Vrålstad, T., Nowak, C., Thines, M., 2018. eDNA-based crayfish plague monitoring is superior to conventional trap-based assessments in year-round detection probability. Hydrobiologia 807, 87–97. https://doi.org/10.1007/s10750-017-3408-8

Zhang, J., You, J., Li, H., Tyler Mehler, W., Zeng, E.Y., 2018. Particle-scale understanding of cypermethrin in sediment: Desorption, bioavailability, and bioaccumulation in benthic invertebrate Lumbriculus variegatus. Science of The Total Environment 642, 638–645. https://doi.org/10.1016/j.scitotenv.2018.06.098

