## Supplemental Figures S1-S3 for "Environmental DNA reveals invasive crayfish microbial associates and ecosystem-wide biodiversity before and after eradication"

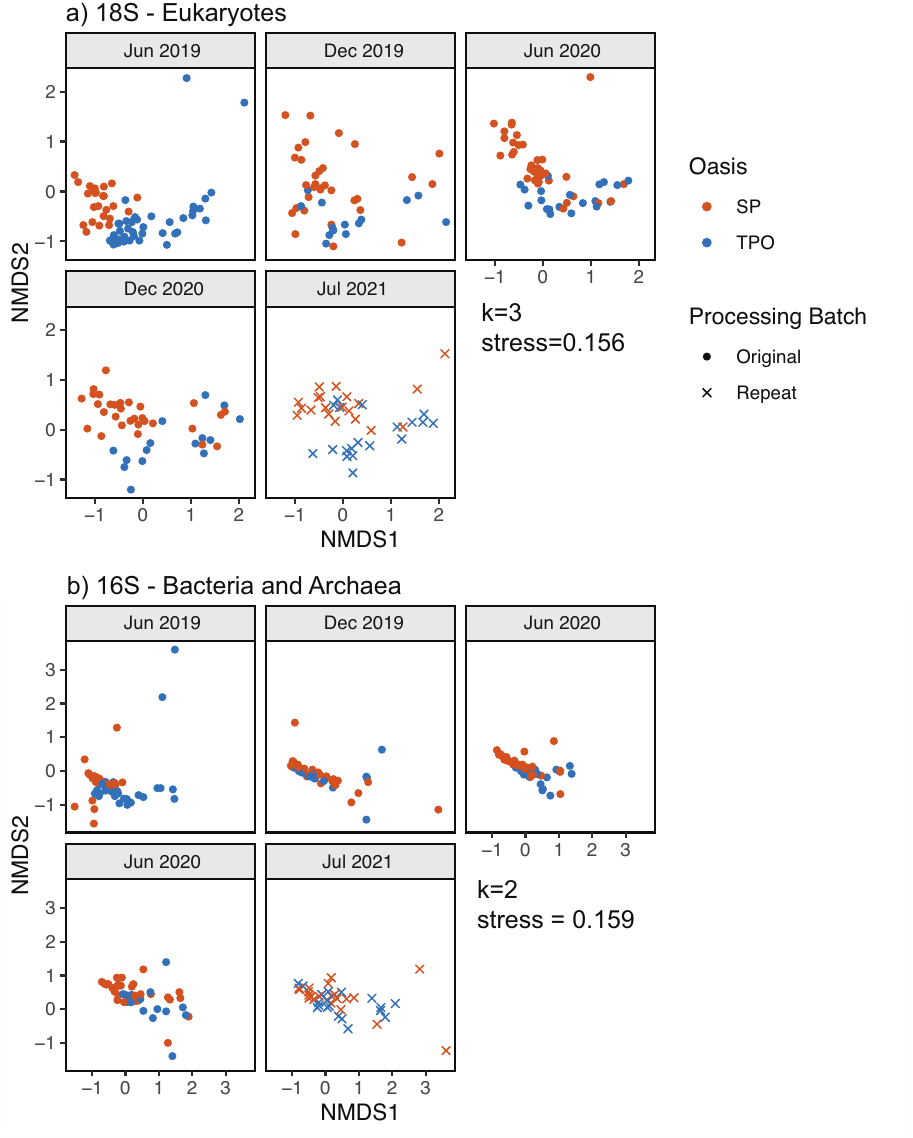

**Supplemental Figure S1.** NMDS plot of a) 18S and b) 16S datasets including repeated samples, faceted by sample processing batch (DNA extraction, library preparation, and sequencing).

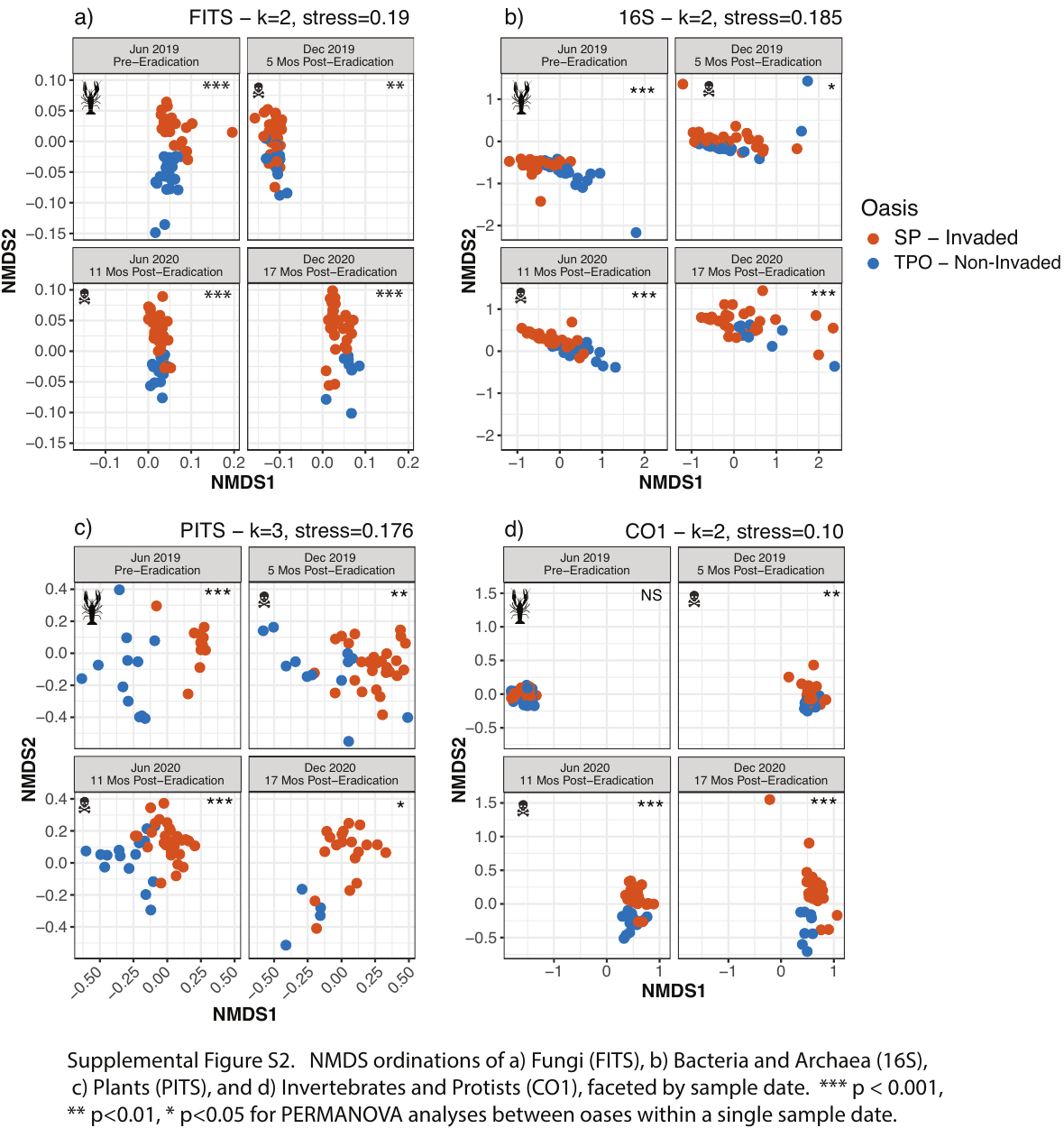

**Supplemental Figure S2.** NMDS ordinations of a) Fungi (FITS), b) Bacteria and Archaea (16S), c) Plants (PITS), and d) Invertebrates and Protists (CO1), faceted by sample date. *** p < 0.001, ** p<0.01, * p<0.05 for PERMANOVA analyses between oases within a single sample date.

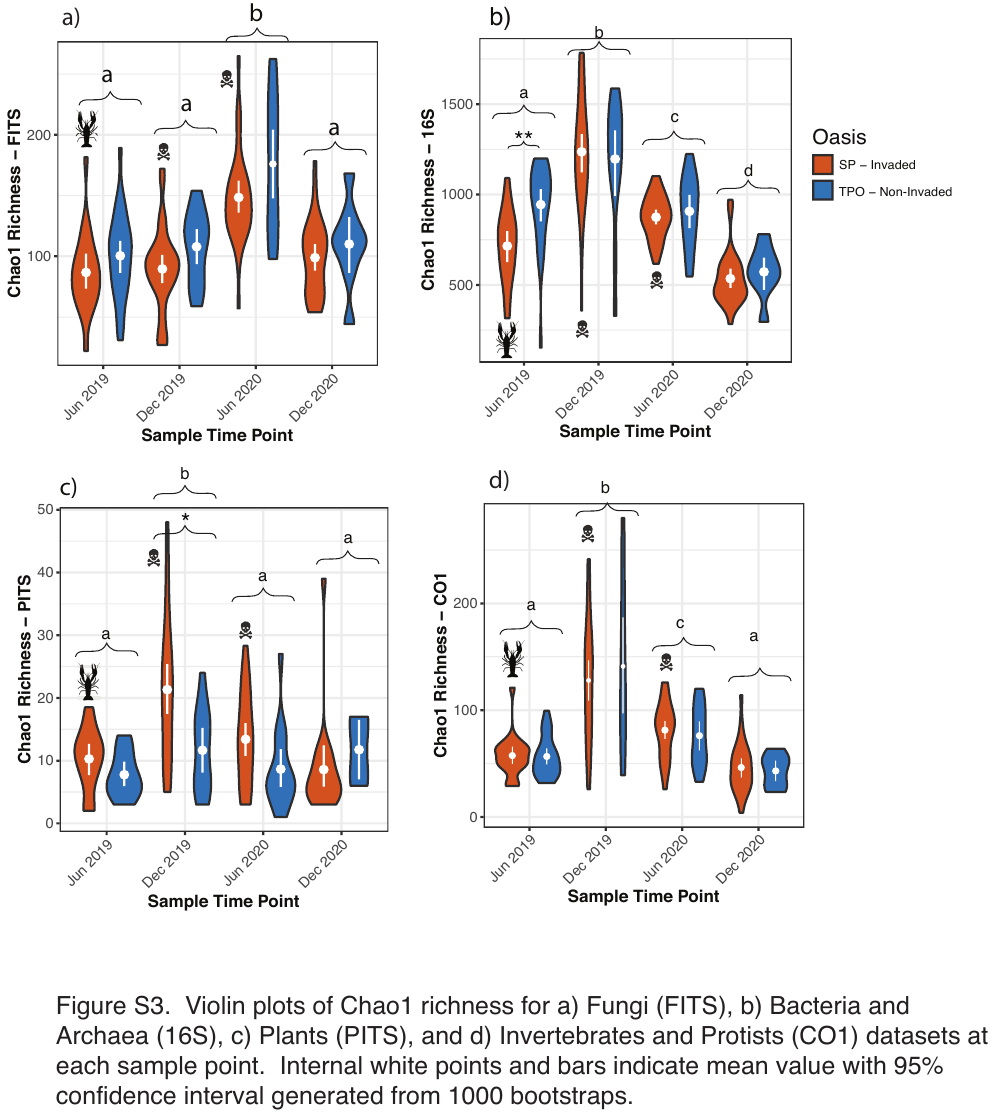

**Supplemental Figure S3**. Violin plots of Chao1 richness for a) Fungi (FITS), b) Bacteria and Archaea (16S), c) Plants (PITS), and d) Invertebrates and Protists (CO1) datasets at each sample point. Internal white points and bars indicate mean value with 95% confidence interval generated from 1000 bootstraps.
