## Supplemental Tables S5 and S6 for "Environmental DNA reveals invasive crayfish microbial associates and ecosystem-wide biodiversity before and after eradication"

**Supplemental Table S5**. Beta Diversity Analysis Results including PERMANOVA with individual sample date included as a stratum, pairwise PERMANOVA comparing Simone Pond (SP) and Thousand Palms Oasis (TPO) communities within a Sample Date, beta dispersion tests with 999 permutations, and post-hoc tukey tests of beta dispersion for sample date, and Oasis*sample date interaction (for brevity, only comparisons within a single date are shown, full results available on request) if significant in the beta dispersion test. Variables that are significant in both PERMANOVA and beta dispersion tests may be showing community differences that are due to differences in dispersion, and not reflecting a true difference in community structure. Variables that are significant in only the PERMANOVA results and not beta dispersion results are showing a true difference in community structure and are bolded in the PERMANOVA results.

| Dataset | Analysis |  |  |  |  | |  |  |
| --- | --- | --- | --- | --- | --- | --- | --- | --- |
| 18S | PERMANOVA | Group | DF | Sum of Squares | R^2^ | | F | P |
|  |  | **Oasis** | **1** | **2.098** | **0.028** | | **5.128** | **<0.001** |
|  |  | Sample Date | 3 | 3.407 | 0.046 | | 2.776 | 0.231 |
|  |  | **Oasis*Sample Date** | **3** | **2.265** | **0.031** | | **1.845** | **<0.001** |
|  |  | Residuals | 162 | 66.273 | 0.895 | |  |  |
|  | Pairwise PERMANOVA | Group (Oases within a Sample Date) | DF | Sum of Squares | R^2^ | | F | P |
|  |  | **Jun 2019 TPO vs SP** | **1** | **1.455** | **0.073** | | **3.637** | **<0.001** |
|  |  | Residuals | 46 | 18.406 | 0.927 | |  |  |
|  |  | **Dec 2019 TPO vs SP** | **1** | **0.739** | **0.042** | | **1.712** | **<0.001** |
|  |  | Residuals | 39 | 16.842 | 0.958 | |  |  |
|  |  | **Jun 2020 TPO vs SP** | **1** | **1.053** | **0.056** | | **2.630** | **<0.001** |
|  |  | Residuals | 44 | 17.607 | 0.944 | |  |  |
|  |  | **Dec 2020 TPO vs SP** | **1** | **0.944** | **0.066** | | **2.323** | **<0.001** |
|  |  | Residuals | 33 | 13.418 | 0.934 | |  |  |
|  | Beta Dispersion Test | Group | DF | Sum of Squares | Mean Square | | F | P |
|  |  | Oasis | 1 | <0.001 | <0.001 | | 0.074 | 0.782 |
|  |  | Residuals | 168 | 0.182 | 0.001 | |  |  |
|  |  | Sample Date | 3 | 0.008 | 0.003 | | 1.831 | 0.142 |
|  |  | Residuals | 166 | 0.247 | 0.001 | |  |  |
|  |  | Oasis*Sample Date | 7 | 0.041 | 0.006 | | 2.458 | 0.017 |
|  |  | Residuals | 162 | 0.390 | 0.002 | |  |  |
|  | Post-Hoc Tukey HSD (beta dispersion) | Group | Pair Compared | Difference | Lower Bound | | Upper Bound | P (adjusted) |
|  |  | Oasis*Sample Date | Jun 2019 TPO vs SP | 0.039 | -0.005 | | 0.082 | 0.129 |
|  |  |  | Dec 2019 TPO vs SP | -0.024 | -0.076 | | 0.028 | 0.846 |
|  |  |  | Jun 2020 TPO vs SP | 0.008 | -0.040 | | 0.055 | 1.000 |
|  |  |  | Dec 2020 TPO vs SP | -0.018 | -0.078 | | 0.043 | 0.986 |
| 16S | PERMANOVA | Group | DF | Sum of Squares | R^2^ | | F | P |
|  |  | **Oasis** | **1** | **1.844** | **0.029** | | **6.296** | **<0.001** |
|  |  | Sample Date | 3 | 11.976 | 0.187 | | 13.628 | <0.001 |
|  |  | **Oasis*Sample Date** | **3** | **2.284** | **0.036** | | **2.599** | **<0.001** |
|  |  | Residuals | 164 | 48.041 | 0.749 | |  |  |
|  | Pairwise PERMANOVAs | Group (Oases within a Sample Date) | DF | Sum of Squares | R^2^ | | F | P |
|  |  | **Jun 2019 TPO vs SP** | **1** | **1.477** | **0.103** | | **5.035** | **0.001** |
|  |  | Residuals | 44 | 12.904 | 0.897 | |  |  |
| 16S | Pairwise PERMANOVAs | Group  (Oases within a Sample Date) | DF | Sum of Squares | R^2^ | | F | P |
|  |  | **Dec 2019 TPO vs SP** | **1** | **0.523** | **0.037** | | **1.630** | **0.018** |
|  |  | Residuals | 43 | 13.805 | 0.963 | |  |  |
|  |  | **Jun 2020 TPO vs SP** | **1** | **1.056** | **0.087** | | **4.219** | **0.001** |
|  |  | Residuals | 44 | 11.016 | 0.913 | |  |  |
|  |  | **Dec 2020 TPO vs SP** | **1** | **0.615** | **0.056** | | **1.969** | **0.009** |
|  |  | Residuals | 33 | 10.316 | 0.944 | |  |  |
|  | Beta Dispersion Test | Group | DF | Sum of Squares | Mean Square | | F | P |
|  |  | Oasis | 1 | <0.001 | <0.001 | | 0.072 | 0.776 |
|  |  | Residuals | 170 | 0.785 | 0.005 | |  |  |
|  |  | Sample Date | 3 | 0.079 | 0.026 | | 3.912 | 0.009 |
|  |  | Residuals | 168 | 1.129 | 0.007 | |  |  |
|  |  | Oasis*Sample Date | 7 | 0.135 | 0.019 | | 2.369 | 0.027 |
|  |  | Residuals | 164 | 1.338 | 0.008 | |  |  |
|  | Post-Hoc Tukey HSD (Beta Dispersion) | Group | Pair Compared | Difference | Lower Bound | | Upper Bound | P (adjusted) |
|  |  | Sample Date | Dec 2019 vs Jun 2019 | 0.004 | -0.041 | | 0.049 | 0.996 |
|  |  |  | Jun 2020 vs Jun 2019 | -0.047 | -0.092 | | -0.003 | 0.032 |
|  |  |  | Dec 2020 vs Jun 2019 | -0.002 | -0.049 | | 0.046 | 1.000 |
|  |  |  | Jun 2020 vs Jun 2019 | -0.051 | -0.096 | | -0.007 | 0.017 |
|  |  |  | Dec 2020 vs Dec 2019 | -0.006 | -0.054 | | 0.042 | 0.990 |
|  |  |  | Dec 2020 vs Jun 2020 | 0.046 | -0.002 | | 0.093 | 0.067 |
|  |  | Oasis*Sample Date | Jun 2019 TPO vs SP | 0.012 | -0.070 | | 0.094 | 1.000 |
|  |  |  | Dec 2019 TPO vs SP | -0.005 | -0.094 | | 0.085 | 1.000 |
|  |  |  | Jun 2020 TPO vs SP | 0.045 | -0.042 | | 0.132 | 0.760 |
|  |  |  | Dec 2020 TPO vs SP | -0.041 | -0.153 | | 0.071 | 0.950 |
| FITS | PERMANOVA | Group | DF | Sum of Squares | R^2^ | | F | P |
|  |  | Oasis | 1 | 1.671 | 0.021 | | 4.121 | <0.001 |
|  |  | Sample Date | 3 | 8.864 | 0.110 | | 7.288 | <0.001 |
|  |  | Oasis*Sample Date | 3 | 3.018 | 0.038 | | 2.481 | <0.001 |
|  |  | Residuals | 165 | 66.893 | 0.832 | |  |  |
|  | Pairwise PERMANOVA | Group (Oases within a Sample Date) | DF | Sum of Squares | R^2^ | | F | P |
|  |  | Jun 2019 TPO vs SP | 1 | 1.918 | 0.096 | | 4.758 | 0.001 |
|  |  | Residuals | 45 | 18.143 | 0.904 | |  |  |
|  |  | Dec 2019 TPO vs SP | 1 | 0.726 | 0.043 | | 1.943 | 0.010 |
|  |  | Residuals | 43 | 16.076 | 0.957 | |  |  |
|  |  | Jun 2020 TPO vs SP | 1 | 1.085 | 0.056 | | 2.587 | 0.001 |
|  |  | Residuals | 44 | 18.459 | 0.944 | |  |  |
|  |  | Dec 2020 TPO vs SP | 1 | 0.843 | 0.056 | | 1.957 | 0.001 |
|  |  | Residuals | 33 | 14.214 | 0.944 | |  |  |
|  | Beta Dispersion Test | Group | DF | Sum of Squares | Mean Square | | F | P |
|  |  | Oasis | 1 | <0.001 | <0.001 | | 0.647 | 0.425 |
|  |  |  | 171 | 0.079 | <0.001 | |  |  |
|  |  | Sample Date | 3 | 0.072 | 0.024 | | 8.700 | 0.001 |
|  |  |  | 169 | 0.466 | 0.003 | |  |  |
|  |  | Oasis*Sample Date | 7 | 0.079 | 0.011 | | 2.605 | 0.020 |
|  |  |  | 165 | 0.713 | 0.004 | |  |  |
|  | Post-Hoc Tukey HSD (beta dispersion) | Group | Pair Compared | Difference | Lower Bound | | Upper Bound | P (adjusted) |
|  |  | Sample Date | Dec 2019 vs Jun 2019 | -0.04616 | -0.075 | | -0.018 | <0.001 |
|  |  |  | Jun 2020 vs Jun 2019 | -0.00158 | -0.030 | | 0.027 | 0.999 |
|  |  |  | Dec 2020 vs Jun 2019 | 0.00277 | -0.028 | | 0.033 | 0.995 |
|  |  |  | Jun 2020 vs Jun 2019 | 0.044582 | 0.016 | | 0.073 | <0.001 |
|  |  |  | Dec 2020 vs Dec 2019 | 0.048929 | 0.018 | | 0.080 | <0.001 |
|  |  |  | Dec 2020 vs Jun 2020 | 0.004347 | -0.026 | | 0.035 | 0.983 |
|  |  |  | Dec 2019 vs Jun 2019 | -0.04616 | -0.075 | | -0.018 | <0.001 |
|  |  | Oasis*Sample Date | Jun 2019 TPO vs SP | 0.041022 | -0.018 | | 0.100 | 0.403 |
|  |  |  | Dec 2019 TPO vs SP | -0.00439 | -0.069 | | 0.061 | 1.000 |
|  |  |  | Jun 2020 TPO vs SP | 0.020003 | -0.043 | | 0.084 | 0.978 |
|  |  |  | Dec 2020 TPO vs SP | -0.02923 | -0.110 | | 0.052 | 0.955 |
| PITS | PERMANOVA | Group | DF | Sum of Squares | R^2^ | | F | P |
|  |  | **Oasis** | **1** | **1.883** | **0.032** | | **4.768** | **<0.001** |
|  |  | **Sample Date** | **3** | **4.883** | **0.084** | | **4.122** | **<0.001** |
|  |  | **Oasis*Sample Date** | **3** | **2.182** | **0.037** | | **1.842** | **<0.001** |
|  |  | Residuals | 125 | 49.361 | 0.847 | |  |  |
|  | Pairwise PERMANOVA | Group (Oases within a Sample Date) | DF | Sum of Squares | R^2^ | | F | P |
|  |  | $Jun 2019 TPO vs SP | 1 | 1.3901 | 0.14372 | | 3.6924 | 0.001 |
|  |  |  | 22 | 8.2825 | 0.85628 | |  |  |
|  |  | **Dec 2019 TPO vs SP** | **1** | **0.966** | **0.05345** | | **2.2024** | **0.001** |
|  |  |  | 39 | 17.107 | 0.94655 | |  |  |
|  |  | **Jun 2020 TPO vs SP** | **1** | **1.0614** | **0.0606** | | **2.7738** | **0.002** |
|  |  |  | 43 | 16.4541 | 0.9394 | |  |  |
|  |  | **Dec 2020 TPO vs SP** | **1** | **0.6627** | **0.08101** | | **1.8512** | **0.035** |
|  |  |  | 21 | 7.5177 | 0.91899 | |  |  |
|  | Beta Dispersion Test | Group | DF | Sum of Squares | Mean Square | | F | P |
|  |  | Oasis | 1 | 0.011 | 0.011 | | 2.402 | 0.134 |
|  |  | Residuals | 131 | 0.594 | 0.005 | |  |  |
|  |  | Sample Date | 3 | 0.096 | 0.032 | | 3.219 | 0.022 |
|  |  | Residuals | 129 | 1.288 | 0.010 | |  |  |
|  |  | Oasis*Sample Date | 7.000 | 0.273 | 0.039 | | 3.830 | 0.002 |
|  |  | Residuals | 125.000 | 1.271 | 0.010 | |  |  |
|  | Post-Hoc Tukey HSD (beta dispersion) | Group | Pair Compared | Difference | Lower Bound | | Upper Bound | P (adjusted) |
|  |  | Sample Date | Dec 2019 vs Jun 2019 | 0.034 | -0.033 | | 0.101 | 0.542 |
|  |  |  | Jun 2020 vs Jun 2019 | -0.011 | -0.077 | | 0.055 | 0.971 |
|  |  |  | Dec 2020 vs Jun 2019 | -0.043 | -0.119 | | 0.033 | 0.454 |
|  |  |  | Jun 2020 vs Jun 2019 | -0.045 | -0.102 | | 0.011 | 0.157 |
|  |  |  | Dec 2020 vs Dec 2019 | -0.077 | -0.145 | | -0.010 | 0.018 |
|  |  |  | Dec 2020 vs Jun 2020 | -0.032 | -0.099 | | 0.035 | 0.597 |
|  |  | Oasis*Sample Date | Jun 2019 TPO vs SP | 0.151 | 0.024 | | 0.278 | 0.008 |
|  |  |  | Dec 2019 TPO vs SP | -0.025 | -0.132 | | 0.082 | 0.996 |
|  |  |  | Jun 2020 TPO vs SP | 0.006 | -0.092 | | 0.105 | 1.000 |
|  |  |  | Dec 2020 TPO vs SP | 0.027 | -0.144 | | 0.198 | 1.000 |
| CO1 | PERMANOVA | Group | DF | Sum of Squares | R^2^ | | F | P |
|  |  | **Oasis** | **1** | **1.333** | **0.017** | | **3.214** | **<0.001** |
|  |  | Sample Date | 3 | 8.333 | 0.108 | | 6.696 | <0.001 |
|  |  | **Oasis*Sample Date** | **3** | **2.546** | **0.033** | | **2.046** | **<0.001** |
|  |  | Residuals | 157 | 65.120 | 0.842 | |  |  |
|  | Pairwise PERMANOVA | Group (Oases within a Sample Date) | DF | Sum of Squares | R^2^ | | F | P (adjusted) |
|  |  | Jun 2019 TPO vs SP | 1 | 1.262 | 0.076 | | 3.359 | 0.001 |
|  |  | Residuals | 41 | 15.409 | 0.924 | |  |  |
|  |  | **Dec 2019 TPO vs SP** | **1** | **0.661** | **0.041** | | **1.662** | **0.004** |
|  |  | Residuals | 39 | 15.508 | 0.959 | |  |  |
|  |  | **Jun 2020 TPO vs SP** | **1** | **0.848** | **0.041** | | **1.899** | **0.001** |
| CO1 |  | Residuals | 44 | 19.645 | 0.959 | |  |  |
|  |  | **Dec 2020 TPO vs SP** | **1** | **0.992** | **0.064** | | **2.249** | **0.001** |
|  |  | Residuals | 33 | 14.558 | 0.936 | |  |  |
|  | Beta Dispersion Test | Group | DF | Sum of Squares | | Mean Square | F | P |
|  |  | Oasis | 1 | 0.001 | 0.001 | | 0.293 | 0.578 |
|  |  |  | 163 | 0.061 | 0.001 | |  |  |
|  |  | Sample Date | 3 | 0.079 | 0.026 | | 12.218 | 0.001 |
|  |  |  | 161 | 0.347 | 0.002 | |  |  |
|  |  | Oasis*Sample Date | 3 | 0.079 | 0.026 | | 12.218 | 0.001 |
|  |  |  | 161 | 0.347 | 0.002 | |  |  |
|  | Post-Hoc Tukey HSD (beta dispersion) | Group | Pair Compared | Difference | Lower Bound | | Upper Bound | P (adjusted) |
|  |  | Sample Date | Dec 2019 vs Jun 2019 | 0.004 | -0.022 | | 0.030 | 0.979 |
|  |  |  | Jun 2020 vs Jun 2019 | 0.046 | 0.021 | | 0.072 | <0.001 |
|  |  |  | Dec 2020 vs Jun 2019 | 0.045 | 0.018 | | 0.072 | <0.001 |
|  |  |  | Jun 2020 vs Jun 2019 | 0.042 | 0.016 | | 0.068 | <0.001 |
|  |  |  | Dec 2020 vs Dec 2019 | 0.041 | 0.013 | | 0.069 | 0.001 |
|  |  |  | Dec 2020 vs Jun 2020 | -0.001 | -0.028 | | 0.026 | 1.000 |
|  |  |  | Dec 2019 vs Jun 2019 | 0.004 | -0.022 | | 0.030 | 0.979 |
|  |  | Oasis*Sample Date | Jun 2019 TPO vs SP | 0.068 | 0.015 | | 0.121 | 0.003 |
|  |  |  | Dec 2019 TPO vs SP | -0.006 | -0.066 | | 0.053 | 1.000 |
|  |  |  | Jun 2020 TPO vs SP | 0.005 | -0.049 | | 0.060 | 1.000 |
|  |  |  | Dec 2020 TPO vs SP | -0.044 | -0.114 | | 0.026 | 0.525 |
| Arthropoda | PERMANOVA | Group | DF | Sum of Squares | R^2^ | | F | P |
|  |  | **Oasis** | **1** | **1.864** | **0.023** | | **4.117** | **<0.001** |
|  |  | **Sample Date** | **3** | **3.497** | **0.043** | | **2.575** | **0.042** |
|  |  | **Oasis*Sample Date** | **3** | **2.569** | **0.032** | | **1.892** | **<0.001** |
|  |  | Residual | 161 | 72.877 | 0.902 | |  |  |
|  | Pairwise PERMANOVA | Group (Oases within a Sample Date) | DF | Sum of Squares | R^2^ | | F | P (adjusted) |
|  |  | **Jun 2019 TPO vs SP** | **1** | **1.285** | **0.059** | | **2.900** | **<0.001** |
|  |  | Residual | 46 | 20.385 | 0.941 | |  |  |
|  |  | **Dec 2019 TPO vs SP** | **1** | **0.871** | **0.046** | | **1.862** | **<0.001** |
|  |  | Residual | 39 | 18.245 | 0.954 | |  |  |
|  |  | **Jun 2020 TPO vs SP** | **1** | **0.879** | **0.041** | | **1.858** | **<0.001** |
|  |  | Residual | 43 | 20.353 | 0.959 | |  |  |
|  |  | **Dec 2020 TPO vs SP** | **1** | **1.309** | **0.086** | | **3.109** | **<0.001** |
|  |  | Residual | 33 | 13.894 | 0.914 | |  |  |
|  | Beta Dispersion Test | Group | DF | Sum of Squares | Mean Square | | F | P |
|  |  | Oasis | 1 | <0.001 | <0.001 | | <0.001 | 0.989 |
|  |  | Residuals | 167 | 0.204 | 0.001 | |  |  |
|  |  | Sample Date | 3 | 0.024 | 0.008 | | 3.544 | 0.016 |
|  |  | Residuals | 165 | 0.376 | 0.002 | |  |  |
|  |  | Oasis*Sample Date | 7 | 0.104 | 0.015 | | 3.582 | 0.002 |
|  |  | Residuals | 161 | 0.668 | 0.004 | |  |  |
|  | Post-Hoc Tukey HSD (beta dispersion) | Group | Pair Compared | Difference | Lower Bound | | Upper Bound | P (adjusted) |
|  |  | Sample Date | Dec 2019 vs Jun 2019 | 0.012 | -0.015 | | 0.038 | 0.652 |
|  |  |  | Jun 2020 vs Jun 2019 | 0.016 | -0.010 | | 0.042 | 0.369 |
|  |  |  | Dec 2020 vs Jun 2019 | -0.016 | -0.044 | | 0.011 | 0.418 |
|  |  |  | Jun 2020 vs Jun 2019 | 0.004 | -0.022 | | 0.031 | 0.976 |
|  |  |  | Dec 2020 vs Dec 2019 | -0.028 | -0.057 | | <0.001 | 0.055 |
|  |  |  | Dec 2020 vs Jun 2020 | -0.032 | -0.060 | | -0.004 | 0.016 |
| Arthropoda |  | Oasis*Sample Date | Jun 2019 TPO vs SP | 0.055 | -0.003 | | 0.112 | 0.077 |
|  |  |  | Dec 2019 TPO vs SP | -0.015 | -0.083 | | 0.053 | 0.997 |
|  |  |  | Jun 2020 TPO vs SP | -0.010 | -0.073 | | 0.052 | 1.000 |
|  |  |  | Dec 2020 TPO vs SP | -0.069 | -0.156 | | 0.017 | 0.221 |

**Supplemental Table S6**. Alpha diversity analysis results including ANOVA and post-hoc tests comparing Chao1 richness between Simone Pond (SP) and Thousand Palms Oasis (TPO), sample dates, and sample date x oasis interaction for each metabarcode dataset. Results significant at the p < 0.05 level are bolded.

| Dataset | Analysis |  |  |  |  |  |  |
| --- | --- | --- | --- | --- | --- | --- | --- |
| 18S | ANOVA | Group | DF | Sum of Squares | Mean Square | F | P |
|  |  | Oasis | **1** | **33264** | **33264** | **6.596** | **0.011** |
|  |  | Sample Date | **3** | **290447** | **96816** | **19.198** | **>0.001** |
|  |  | Oasis*Sample Date | 3 | 36392 | 12131 | 2.405 | 0.069 |
|  |  | Residuals | 162 | 816974 | 5043 |  |  |
|  | Post-Hoc Tukey HSD | Group | Pair Compared | Difference | Lower Bound | Upper Bound | P (adjusted) |
|  |  | Oasis | **TPO*SP** | **29.061** | **6.716** | **51.405** | **0.011** |
|  |  | Sample Date | Dec 2019-Jun 2019 | -33.568 | -72.771 | 5.635 | 0.121 |
|  |  |  | **Jun 2020-Jun 2019** | **-100.190** | **-138.226** | **-62.154** | **>0.001** |
|  |  |  | **Dec 2020-Jun 2019** | **-80.293** | **-121.268** | **-39.318** | **>0.001** |
|  |  |  | **Jun 2020-Dec 2019** | **-66.622** | **-106.215** | **-27.028** | **>0.001** |
|  |  |  | **Dec 2020-Dec 2019** | **-46.724** | **-89.149** | **-4.300** | **0.025** |
|  |  |  | Dec 2020-Jun 2020 | 19.898 | -21.451 | 61.246 | 0.597 |
|  |  |  | Dec 2019-Jun 2019 | -33.568 | -72.771 | 5.635 | 0.121 |
|  |  |  | **Jun 2020-Jun 2019** | **-100.190** | **-138.226** | **-62.154** | **>0.001** |
|  |  | Oasis*Sample Date | SP*Dec 2019-SP*Jun 2019 | -34.904 | -97.392 | 27.584 | 0.677 |
|  |  |  | **SP*Dec 2019-TPO*Jun 2019** | **-69.029** | **-127.352** | **-10.707** | **0.009** |
|  |  |  | **SP*Jun 2020-SP*Jun 2019** | **-83.852** | **-145.487** | **-22.216** | **0.001** |
|  |  |  | SP*Jun 2020-SP*Dec 2019 | -48.947 | -105.287 | 7.393 | 0.140 |
|  |  |  | **SP*Jun 2020-TPO*Jun 2019** | **-117.977** | **-175.385** | **-60.569** | **>0.001** |
|  |  |  | **SP*Jun 2020-TPO*Dec 2019** | **-92.306** | **-166.452** | **-18.161** | **0.005** |
|  |  |  | SP*Dec 2020-SP*Jun 2019 | -61.789 | -125.241 | 1.664 | 0.062 |
|  |  |  | SP*Dec 2020-SP*Dec 2019 | -26.884 | -85.207 | 31.438 | 0.849 |
|  |  |  | SP*Dec 2020-SP*Jun 2020 | 22.063 | -35.345 | 79.471 | 0.937 |
|  |  |  | **SP*Dec 2020-TPO*Jun 2019** | **-95.914** | **-155.268** | **-36.559** | **>0.001** |
|  |  |  | SP*Dec 2020-TPO*Dec 2019 | -70.243 | -145.906 | 5.419 | 0.090 |
|  |  |  | SP*Dec 2020-TPO*Jun 2020 | 34.371 | -35.859 | 104.600 | 0.805 |
|  |  |  | TPO*Jun 2019-SP*Jun 2019 | 34.125 | -29.328 | 97.578 | 0.718 |
|  |  |  | TPO*Dec 2019-SP*Jun 2019 | 8.455 | -70.464 | 87.373 | 1.000 |
|  |  |  | TPO*Dec 2019-SP*Dec 2019 | 43.359 | -31.496 | 118.214 | 0.635 |
|  |  |  | TPO*Dec 2019-TPO*Jun 2019 | -25.670 | -101.333 | 49.992 | 0.967 |
|  |  |  | **TPO*Jun 2020-SP*Jun 2019** | **-96.159** | **-169.885** | **-22.434** | **0.002** |
|  |  |  | TPO*Jun 2020-SP*Dec 2019 | -61.255 | -130.614 | 8.104 | 0.126 |
|  |  |  | TPO*Jun 2020-SP*Jun 2020 | -12.307 | -80.900 | 56.285 | 0.999 |
|  |  |  | **TPO*Jun 2020-TPO*Jun 2019** | **-130.284** | **-200.514** | **-60.055** | **>0.001** |
|  |  |  | **TPO*Jun 2020-TPO*Dec 2019** | **-104.614** | **-189.077** | **-20.151** | **0.005** |
|  |  |  | **TPO*Dec 2020-SP*Jun 2019** | **-101.220** | **-191.827** | **-10.612** | **0.017** |
|  |  |  | TPO*Dec 2020-SP*Dec 2019 | -66.315 | -153.407 | 20.777 | 0.280 |
|  |  |  | TPO*Dec 2020-SP*Jun 2020 | -17.368 | -103.850 | 69.115 | 0.999 |
|  |  |  | TPO*Dec 2020-SP*Dec 2020 | -39.431 | -127.218 | 48.356 | 0.865 |
|  |  |  | TPO*Dec 2020-TPO*Jun 2019 | -135.345 | -223.131 | -47.558 | >0.001 |
|  |  |  | TPO*Dec 2020-TPO*Dec 2019 | -109.674 | -209.215 | -10.134 | 0.020 |
|  |  |  | TPO*Dec 2020-TPO*Jun 2020 | -5.060 | -100.536 | 90.416 | 1.000 |
| FITS | ANOVA | Group | DF | Sum of Squares | Mean Square | F | P |
|  |  | Oasis | **1** | **7184** | **7184** | **5.368** | **0.0217** |
|  |  | Sample Date | **3** | **131305** | **43768** | **32.709** | **<0.001** |
|  |  | Oasis*Sample Date | 3 | 1448 | 483 | 0.361 | 0.7815 |
|  |  | Residuals | 165 | 220792 | 1338 |  |  |
|  | Post-Hoc Tukey HSD | Group | Pair Compared | Difference | Lower Bound | Upper Bound | P (adjusted) |
|  |  | Oasis | **TPO-SP** | **13.391** | **1.980** | **24.803** | **0.022** |
|  |  | Sample Date | Dec 2019-Jun 2019 | 4.304 | -15.497 | 24.106 | 0.943 |
|  |  |  | **Jun 2020-Jun 2019** | **66.202** | **46.511** | **85.893** | **>0.001** |
|  |  |  | Dec 2020-Jun 2019 | 11.470 | -9.727 | 32.667 | 0.498 |
|  |  |  | **Jun 2020-Dec 2019** | **61.897** | **41.991** | **81.804** | **>0.001** |
|  |  |  | Dec 2020-Dec 2019 | 7.166 | -14.232 | 28.563 | 0.821 |
|  |  |  | **Dec 2020-Jun 2020** | **-54.732** | **-76.027** | **-33.436** | **>0.001** |
|  |  | Oasis*Sample Date | TPO*Jun 2019-SP*Jun 2019 | 13.642 | -19.309 | 46.594 | 0.908 |
|  |  |  | SP*Dec 2019-SP*Jun 2019 | 2.906 | -28.835 | 34.648 | 1.000 |
|  |  |  | TPO*Dec 2019-SP*Jun 2019 | 21.237 | -17.514 | 59.988 | 0.698 |
| FITS | Post-Hoc Tukey HSD | Group | Pair Compared | Difference | Lower Bound | Upper Bound | P (adjusted) |
|  |  |  | **SP*Jun 2020-SP*Jun 2019** | **61.697** | **29.955** | **93.438** | **>0.001** |
|  |  |  | **TPO*Jun 2020-SP*Jun 2019** | **89.329** | **51.361** | **127.297** | **>0.001** |
|  |  |  | SP*Dec 2020-SP*Jun 2019 | 12.122 | -20.556 | 44.800 | 0.947 |
|  |  |  | TPO*Dec 2020-SP*Jun 2019 | 23.268 | -23.394 | 69.930 | 0.790 |
|  |  |  | SP*Dec 2019-TPO*Jun 2019 | -10.736 | -40.603 | 19.131 | 0.955 |
|  |  |  | TPO*Dec 2019-TPO*Jun 2019 | 7.594 | -29.636 | 44.825 | 0.998 |
|  |  |  | **SP*Jun 2020-TPO*Jun 2019** | **48.054** | **18.187** | **77.921** | **>0.001** |
|  |  |  | **TPO*Jun 2020-TPO*Jun 2019** | **75.687** | **39.272** | **112.102** | **>0.001** |
|  |  |  | SP*Dec 2020-TPO*Jun 2019 | -1.520 | -32.380 | 29.339 | 1.000 |
|  |  |  | TPO*Dec 2020-TPO*Jun 2019 | 9.626 | -35.782 | 55.033 | 0.998 |
|  |  |  | TPO*Dec 2019-SP*Dec 2019 | 18.330 | -17.834 | 54.495 | 0.775 |
|  |  |  | **SP*Jun 2020-SP*Dec 2019** | **58.790** | **30.263** | **87.317** | **>0.001** |
|  |  |  | **TPO*Jun 2020-SP*Dec 2019** | **86.423** | **51.099** | **121.747** | **>0.001** |
|  |  |  | SP*Dec 2020-SP*Dec 2019 | 9.215 | -20.349 | 38.780 | 0.980 |
|  |  |  | TPO*Dec 2020-SP*Dec 2019 | 20.362 | -24.176 | 64.899 | 0.854 |
|  |  |  | **SP*Jun 2020-TPO*Dec 2019** | **40.460** | **4.295** | **76.624** | **0.017** |
|  |  |  | **TPO*Jun 2020-TPO*Dec 2019** | **68.092** | **26.357** | **109.828** | **>0.001** |
|  |  |  | SP*Dec 2020-TPO*Dec 2019 | -9.115 | -46.103 | 27.874 | 0.995 |
|  |  |  | TPO*Dec 2020-TPO*Dec 2019 | 2.031 | -47.745 | 51.808 | 1.000 |
|  |  |  | TPO*Jun 2020-SP*Jun 2020 | 27.633 | -7.691 | 62.957 | 0.248 |
|  |  |  | **SP*Dec 2020-SP*Jun 2020** | **-49.575** | **-79.139** | **-20.010** | **>0.001** |
|  |  |  | TPO*Dec 2020-SP*Jun 2020 | -38.428 | -82.966 | 6.109 | 0.146 |
|  |  |  | **SP*Dec 2020-TPO*Jun 2020** | **-77.207** | **-113.375** | **-41.040** | **>0.001** |
|  |  |  | **TPO*Dec 2020-TPO*Jun 2020** | **-66.061** | **-115.230** | **-16.892** | **0.001** |
|  |  |  | TPO*Dec 2020-SP*Dec 2020 | 11.146 | -34.063 | 56.356 | 0.995 |
| 16S | ANOVA | Group | DF | Sum of Squares | Mean Square | F | P |
|  |  | Oasis | 1 | 88198 | 88198 | 2.726 | 0.10062 |
|  |  | Sample Date | 3 | 4931519 | 1643840 | 50.813 | <0.001 |
|  |  | Oasis*Sample Date | 3 | 442461 | 147487 | 4.559 | 0.00427 |
|  |  | Residuals | 164 | 5305552 | 32351 |  |  |
|  | Post-Hoc Tukey HSD | Group | Pair Compared | Difference | Lower Bound | Upper Bound | P (adjusted) |
|  |  | Oasis | TPO-SP | 47.163 | -9.237 | 103.563 | 0.101 |
|  |  | Sample Date | **Dec 2019-Jun 2019** | **310.214** | **212.330** | **408.098** | **>0.001** |
|  |  |  | **Jun 2020-Jun 2019** | **118.449** | **21.104** | **215.793** | **0.010** |
|  |  |  | **Dec 2020-Jun 2019** | **-169.826** | **-274.540** | **-65.112** | **>0.001** |
|  |  |  | **Jun 2020-Dec 2019** | **-191.766** | **-289.650** | **-93.882** | **>0.001** |
|  |  |  | **Dec 2020-Dec 2019** | **-480.041** | **-585.256** | **-374.825** | **>0.001** |
|  |  |  | **Dec 2020-Jun 2020** | **-288.275** | **-392.989** | **-183.561** | **>0.001** |
|  |  | Oasis*Sample Date | **TPO*Jun 2019-SP*Jun 2019** | **208.712** | **45.239** | **372.186** | **0.003** |
|  |  |  | **SP*Dec 2019-SP*Jun 2019** | **431.694** | **275.610** | **587.779** | **>0.001** |
|  |  |  | **TPO*Dec 2019-SP*Jun 2019** | **370.595** | **180.045** | **561.145** | **>0.001** |
|  |  |  | **SP*Jun 2020-SP*Jun 2019** | **222.469** | **66.384** | **378.553** | **0.001** |
|  |  |  | **TPO*Jun 2020-SP*Jun 2019** | **219.886** | **33.186** | **406.586** | **0.009** |
|  |  |  | SP*Dec 2020-SP*Jun 2019 | -74.455 | -235.141 | 86.231 | 0.846 |
|  |  |  | TPO*Dec 2020-SP*Jun 2019 | -60.423 | -289.875 | 169.030 | 0.992 |
|  |  |  | **SP*Dec 2019-TPO*Jun 2019** | **222.982** | **74.528** | **371.436** | **>0.001** |
|  |  |  | TPO*Dec 2019-TPO*Jun 2019 | 161.883 | -22.469 | 346.234 | 0.131 |
|  |  |  | SP*Jun 2020-TPO*Jun 2019 | 13.756 | -134.698 | 162.210 | 1.000 |
|  |  |  | TPO*Jun 2020-TPO*Jun 2019 | 11.173 | -169.196 | 191.543 | 1.000 |
|  |  |  | **SP*Dec 2020-TPO*Jun 2019** | **-283.167** | **-436.452** | **-129.883** | **>0.001** |
|  |  |  | **TPO*Dec 2020-TPO*Jun 2019** | **-269.135** | **-493.467** | **-44.803** | **0.007** |
|  |  |  | TPO*Dec 2019-SP*Dec 2019 | -61.099 | -238.931 | 116.733 | 0.965 |
|  |  |  | **SP*Jun 2020-SP*Dec 2019** | **-209.226** | **-349.502** | **-68.950** | **>0.001** |
|  |  |  | **TPO*Jun 2020-SP*Dec 2019** | **-211.809** | **-385.509** | **-38.108** | **0.006** |
|  |  | Oasis*Sample Date | **SP*Dec 2020-SP*Dec 2019** | **-506.149** | **-651.528** | **-360.771** | **>0.001** |
|  |  |  | **TPO*Dec 2020-SP*Dec 2019** | **-492.117** | **-711.122** | **-273.112** | **>0.001** |
|  |  |  | SP*Jun 2020-TPO*Dec 2019 | -148.127 | -325.959 | 29.705 | 0.179 |
|  |  |  | TPO*Jun 2020-TPO*Dec 2019 | -150.710 | -355.938 | 54.519 | 0.325 |
|  |  |  | **SP*Dec 2020-TPO*Dec 2019** | **-445.050** | **-626.934** | **-263.166** | **>0.001** |
|  |  |  | **TPO*Dec 2020-TPO*Dec 2019** | **-431.018** | **-675.784** | **-186.252** | **>0.001** |
| 16S | Post-Hoc Tukey HSD | Group | Pair Compared | Difference | Lower Bound | Upper Bound | P (adjusted) |
|  |  | Oasis*Sample Date | TPO*Jun 2020-SP*Jun 2020 | -2.583 | -176.283 | 171.118 | 1.000 |
|  |  |  | **SP*Dec 2020-SP*Jun 2020** | **-296.924** | **-442.302** | **-151.545** | **>0.001** |
|  |  |  | **TPO*Dec 2020-SP*Jun 2020** | **-282.891** | **-501.897** | **-63.886** | **0.003** |
|  |  |  | **SP*Dec 2020-TPO*Jun 2020** | **-294.341** | **-472.187** | **-116.494** | **>0.001** |
|  |  |  | **TPO*Dec 2020-TPO*Jun 2020** | **-280.308** | **-522.089** | **-38.528** | **0.011** |
|  |  |  | TPO*Dec 2020-SP*Dec 2020 | 14.032 | -208.276 | 236.341 | 1.000 |
| PITS | ANOVA | Group | DF | Sum of Squares | Mean Square | F | P |
|  |  | Oasis | **1** | **716** | **716.2** | **11.24** | **0.001** |
|  |  | Sample Date | **3** | **1964** | **654.5** | **10.27** | **>0.001** |
|  |  | Oasis*Sample Date | 3 | 443 | 147.8 | 2.32 | 0.07854 |
|  |  | Residuals | 126 | 8028 | 63.7 |  |  |
|  | Post-Hoc Tukey HSD | Group | Pair Compared | Difference | Lower Bound | Upper Bound | P (adjusted) |
|  |  | Oasis | **TPO-SP** | **-4.952** | **-7.875** | **-2.029** | **0.001** |
|  |  | Sample Date | **Dec 2019-Jun 2019** | **8.502** | **3.228** | **13.776** | **>0.001** |
|  |  |  | Jun 2020-Jun 2019 | 2.028 | -3.156 | 7.213 | 0.739 |
|  |  |  | Dec 2020-Jun 2019 | -1.445 | -7.450 | 4.559 | 0.923 |
|  |  |  | **Jun 2020-Dec 2019** | **-6.474** | **-10.961** | **-1.987** | **0.001** |
|  |  |  | **Dec 2020-Dec 2019** | **-9.948** | **-15.362** | **-4.533** | **>0.001** |
|  |  |  | Dec 2020-Jun 2020 | -3.474 | -8.801 | 1.853 | 0.329 |
|  |  | Oasis*Sample Date | **SP*Dec 2019-SP*Jun 2019** | **11.050** | **2.837** | **19.262** | **0.002** |
|  |  |  | **SP*Dec 2019-TPO*Jun 2019** | **13.569** | **5.123** | **22.015** | **>0.001** |
|  |  |  | SP*Jun 2020-SP*Jun 2019 | 3.136 | -5.034 | 11.307 | 0.935 |
|  |  |  | **SP*Jun 2020-SP*Dec 2019** | **-7.913** | **-14.321** | **-1.506** | **0.005** |
|  |  |  | SP*Jun 2020-TPO*Jun 2019 | 5.656 | -2.749 | 14.060 | 0.437 |
|  |  |  | SP*Jun 2020-TPO*Dec 2019 | 1.753 | -6.652 | 10.157 | 0.998 |
|  |  |  | SP*Dec 2020-SP*Jun 2019 | -1.690 | -10.547 | 7.166 | 0.999 |
|  |  |  | **SP*Dec 2020-SP*Dec 2019** | **-12.740** | **-20.002** | **-5.478** | **>0.001** |
|  |  |  | SP*Dec 2020-SP*Jun 2020 | -4.827 | -12.041 | 2.388 | 0.445 |
|  |  |  | SP*Dec 2020-TPO*Jun 2019 | 0.829 | -8.244 | 9.902 | 1.000 |
|  |  |  | SP*Dec 2020-TPO*Dec 2019 | -3.074 | -12.147 | 5.999 | 0.967 |
|  |  |  | SP*Dec 2020-TPO*Jun 2020 | -0.054 | -8.553 | 8.444 | 1.000 |
|  |  |  | TPO*Jun 2019-SP*Jun 2019 | -2.519 | -12.369 | 7.331 | 0.993 |
|  |  |  | TPO*Dec 2019-SP*Jun 2019 | 1.384 | -8.466 | 11.234 | 1.000 |
|  |  |  | **TPO*Dec 2019-SP*Dec 2019** | **-9.666** | **-18.112** | **-1.221** | **0.013** |
|  |  |  | TPO*Dec 2019-TPO*Jun 2019 | 3.903 | -6.142 | 13.948 | 0.931 |
|  |  |  | TPO*Jun 2020-SP*Jun 2019 | -1.636 | -10.960 | 7.688 | 0.999 |
|  |  |  | **TPO*Jun 2020-SP*Dec 2019** | **-12.686** | **-20.511** | **-4.860** | **>0.001** |
|  |  |  | TPO*Jun 2020-SP*Jun 2020 | -4.772 | -12.553 | 3.009 | 0.560 |
|  |  |  | TPO*Jun 2020-TPO*Jun 2019 | 0.883 | -8.646 | 10.413 | 1.000 |
|  |  |  | TPO*Jun 2020-TPO*Dec 2019 | -3.019 | -12.549 | 6.510 | 0.977 |
|  |  |  | TPO*Dec 2020-SP*Jun 2019 | 1.481 | -12.588 | 15.549 | 1.000 |
|  |  |  | TPO*Dec 2020-SP*Dec 2019 | -9.569 | -22.693 | 3.555 | 0.331 |
|  |  |  | TPO*Dec 2020-SP*Jun 2020 | -1.656 | -14.753 | 11.442 | 1.000 |
|  |  |  | TPO*Dec 2020-SP*Dec 2020 | 3.171 | -10.365 | 16.707 | 0.996 |
|  |  |  | TPO*Dec 2020-TPO*Jun 2019 | 4.000 | -10.206 | 18.206 | 0.988 |
|  |  |  | TPO*Dec 2020-TPO*Dec 2019 | 0.097 | -14.109 | 14.303 | 1.000 |
|  |  |  | TPO*Dec 2020-TPO*Jun 2020 | 3.117 | -10.730 | 16.963 | 0.997 |
| CO1 | ANOVA | Group | DF | Sum of Squares | Mean Square | F | P |
|  |  | Oasis | 1 | 435 | 435 | 0.304 | 0.582 |
|  |  | **Sample Date** | **3** | **173587** | **57862** | **40.434** | **<0.001** |
|  |  | Oasis*Sample Date | 3 | 1737 | 579 | 0.405 | 0.75 |
|  |  | Residuals | 157 | 224669 | 1431 |  |  |
|  | Post-Hoc Tukey HSD | Group | Pair Compared | Difference | Lower Bound | Upper Bound | P (adjusted) |
|  |  | Oasis | TPO-SP | -3.401 | -15.584 | 8.782 | 0.582 |
|  |  | Sample Date | **Dec 2019-Jun 2019** | **73.927** | **52.485** | **95.369** | **>0.001** |
|  |  |  | **Jun 2020-Jun 2019** | **21.914** | **1.077** | **42.751** | **0.035** |
|  |  |  | Dec 2020-Jun 2019 | -12.471 | -34.834 | 9.892 | 0.471 |
| CO1 | Post-Hoc Tukey HSD | Group | Pair Compared | Difference | Lower Bound | Upper Bound | P (adjusted) |
|  |  | Sample Date | **Jun 2020-Dec 2019** | **-52.013** | **-73.111** | **-30.915** | **>0.001** |
|  |  |  | **Dec 2020-Dec 2019** | **-86.398** | **-109.005** | **-63.791** | **>0.001** |
|  |  |  | **Dec 2020-Jun 2020** | **-34.385** | **-56.419** | **-12.352** | **>0.001** |
|  |  | Oasis*Sample Date | **SP*Dec 2019-SP*Jun 2019** | **70.523** | **36.743** | **104.304** | **>0.001** |
|  |  |  | **SP*Dec 2019-TPO*Jun 2019** | **71.336** | **38.886** | **103.787** | **>0.001** |
|  |  |  | SP*Jun 2020-SP*Jun 2019 | 23.868 | -9.464 | 57.201 | 0.358 |
|  |  |  | **SP*Jun 2020-SP*Dec 2019** | **-46.655** | **-76.680** | **-16.631** | **>0.001** |
|  |  |  | SP*Jun 2020-TPO*Jun 2019 | 24.681 | -7.303 | 56.665 | 0.263 |
|  |  |  | **SP*Jun 2020-TPO*Dec 2019** | **-59.613** | **-99.126** | **-20.100** | **>0.001** |
|  |  |  | SP*Dec 2020-SP*Jun 2019 | -11.139 | -45.426 | 23.148 | 0.974 |
|  |  |  | **SP*Dec 2020-SP*Dec 2019** | **-81.663** | **-112.743** | **-50.582** | **>0.001** |
|  |  |  | SP*Dec 2020-SP*Jun 2020 | -35.007 | -65.601 | -4.414 | 0.013 |
|  |  |  | SP*Dec 2020-TPO*Jun 2019 | -10.326 | -43.304 | 22.651 | 0.979 |
|  |  |  | **SP*Dec 2020-TPO*Dec 2019** | **-94.620** | **-134.942** | **-54.299** | **>0.001** |
|  |  |  | SP*Dec 2020-TPO*Jun 2020 | -29.859 | -67.285 | 7.567 | 0.225 |
|  |  |  | TPO*Jun 2019-SP*Jun 2019 | -0.813 | -36.346 | 34.720 | 1.000 |
|  |  |  | **TPO*Dec 2019-SP*Jun 2019** | **83.481** | **41.044** | **125.918** | **>0.001** |
|  |  |  | TPO*Dec 2019-SP*Dec 2019 | 12.958 | -26.934 | 52.849 | 0.974 |
|  |  |  | **TPO*Dec 2019-TPO*Jun 2019** | **84.294** | **42.908** | **125.680** | **>0.001** |
|  |  |  | TPO*Jun 2020-SP*Jun 2019 | 18.720 | -20.976 | 58.416 | 0.833 |
|  |  |  | **TPO*Jun 2020-SP*Dec 2019** | **-51.803** | **-88.766** | **-14.841** | **0.001** |
|  |  |  | TPO*Jun 2020-SP*Jun 2020 | -5.148 | -41.702 | 31.405 | 1.000 |
|  |  |  | TPO*Jun 2020-TPO*Jun 2019 | 19.533 | -19.038 | 58.104 | 0.776 |
|  |  |  | **TPO*Jun 2020-TPO*Dec 2019** | **-64.761** | **-109.773** | **-19.750** | **>0.001** |
|  |  |  | TPO*Dec 2020-SP*Jun 2019 | -14.311 | -62.929 | 34.307 | 0.985 |
|  |  |  | **TPO*Dec 2020-SP*Dec 2019** | **-84.834** | **-131.246** | **-38.421** | **>0.001** |
|  |  |  | TPO*Dec 2020-SP*Jun 2020 | -38.179 | -84.266 | 7.909 | 0.185 |
|  |  |  | TPO*Dec 2020-SP*Dec 2020 | -3.171 | -49.954 | 43.611 | 1.000 |
|  |  |  | TPO*Dec 2020-TPO*Jun 2019 | -13.498 | -61.201 | 34.206 | 0.988 |
|  |  |  | **TPO*Dec 2020-TPO*Dec 2019** | **-97.792** | **-150.838** | **-44.745** | **>0.001** |
|  |  |  | TPO*Dec 2020-TPO*Jun 2020 | -33.031 | -83.911 | 17.850 | 0.489 |
| Arthropoda | ANOVA | Group | DF | Sum of Squares | Mean Square | F | P |
|  |  | Oasis | **1** | **249** | **248.6** | **8.333** | **0.004** |
|  |  | Sample Date | **3** | **955** | **318.3** | **10.67** | **>0.001** |
|  |  | Oasis*Sample Date | **3** | **967** | **322.4** | **10.806** | **>0.001** |
|  |  | Residuals | 161 | 4803 | 29.8 |  |  |
|  | Post-Hoc Tukey HSD | Group | Pair Compared | Difference | Lower Bound | Upper Bound | P (adjusted) |
|  |  | Oasis | **TPO-SP** | **2.517** | **0.795** | **4.238** | **0.004** |
|  |  | Sample Date | Dec 2019-Jun 2019 | -2.572 | -5.588 | 0.443 | 0.124 |
|  |  |  | **Jun 2020-Jun 2019** | **-5.206** | **-8.148** | **-2.263** | **>0.001** |
|  |  |  | **Dec 2020-Jun 2019** | **-5.680** | **-8.832** | **-2.528** | **>0.001** |
|  |  |  | Jun 2020-Dec 2019 | -2.633 | -5.695 | 0.428 | 0.119 |
|  |  |  | Dec 2020-Dec 2019 | -3.108 | -6.371 | 0.156 | 0.068 |
|  |  |  | Dec 2020-Jun 2020 | -0.474 | -3.670 | 2.721 | 0.980 |
|  |  | Oasis*Sample Date | **SP*Dec 2019-SP*Jun 2019** | **-8.496** | **-13.303** | **-3.689** | **>0.001** |
|  |  |  | SP*Dec 2019-TPO*Jun 2019 | -3.790 | -8.276 | 0.696 | 0.166 |
|  |  |  | **SP*Jun 2020-SP*Jun 2019** | **-8.563** | **-13.336** | **-3.790** | **>0.001** |
|  |  |  | SP*Jun 2020-SP*Dec 2019 | -0.067 | -4.435 | 4.302 | 1.000 |
|  |  |  | SP*Jun 2020-TPO*Jun 2019 | -3.857 | -8.307 | 0.593 | 0.142 |
|  |  |  | **SP*Jun 2020-TPO*Dec 2019** | **-8.942** | **-14.671** | **-3.212** | **>0.001** |
|  |  |  | **SP*Dec 2020-SP*Jun 2019** | **-10.089** | **-14.969** | **-5.208** | **>0.001** |
|  |  |  | SP*Dec 2020-SP*Dec 2019 | -1.593 | -6.079 | 2.894 | 0.958 |
|  |  |  | SP*Dec 2020-SP*Jun 2020 | -1.526 | -5.976 | 2.924 | 0.965 |
|  |  |  | **SP*Dec 2020-TPO*Jun 2019** | **-5.383** | **-9.948** | **-0.817** | **0.009** |
|  |  |  | **SP*Dec 2020-TPO*Dec 2019** | **-10.468** | **-16.288** | **-4.648** | **>0.001** |
|  |  |  | SP*Dec 2020-TPO*Jun 2020 | -1.926 | -7.328 | 3.476 | 0.957 |
|  |  |  | TPO*Jun 2019-SP*Jun 2019 | -4.706 | -9.587 | 0.175 | 0.068 |
|  |  |  | TPO*Dec 2019-SP*Jun 2019 | 0.379 | -5.692 | 6.449 | 1.000 |
|  |  |  | **TPO*Dec 2019-SP*Dec 2019** | **8.875** | **3.117** | **14.633** | **>0.001** |
|  |  |  | TPO*Dec 2019-TPO*Jun 2019 | 5.085 | -0.735 | 10.905 | 0.135 |
| Arthropoda | Post-Hoc Tukey HSD | Group | Pair Compared | Difference | Lower Bound | Upper Bound | P (adjusted) |
|  |  | Oasis*Sample Date | **TPO*Jun 2020-SP*Jun 2019** | **-8.163** | **-13.834** | **-2.492** | **>0.001** |
|  |  |  | TPO*Jun 2020-SP*Dec 2019 | 0.333 | -5.002 | 5.669 | 1.000 |
|  |  |  | TPO*Jun 2020-SP*Jun 2020 | 0.400 | -4.905 | 5.705 | 1.000 |
|  |  |  | TPO*Jun 2020-TPO*Jun 2019 | -3.457 | -8.859 | 1.945 | 0.509 |
|  |  |  | **TPO*Jun 2020-TPO*Dec 2019** | **-8.542** | **-15.039** | **-2.045** | **0.002** |
|  |  |  | TPO*Dec 2020-SP*Jun 2019 | -6.059 | -13.028 | 0.911 | 0.140 |
|  |  |  | TPO*Dec 2020-SP*Dec 2019 | 2.438 | -4.262 | 9.137 | 0.952 |
|  |  |  | TPO*Dec 2020-SP*Jun 2020 | 2.504 | -4.171 | 9.179 | 0.944 |
|  |  |  | TPO*Dec 2020-SP*Dec 2020 | 4.030 | -2.723 | 10.783 | 0.599 |
|  |  |  | TPO*Dec 2020-TPO*Jun 2019 | -1.353 | -8.105 | 5.400 | 0.999 |
|  |  |  | TPO*Dec 2020-TPO*Dec 2019 | -6.438 | -14.094 | 1.219 | 0.170 |
|  |  |  | TPO*Dec 2020-TPO*Jun 2020 | 2.104 | -5.240 | 9.448 | 0.987 |
